# Winter is coming: pathogen emergence in seasonal environments

**DOI:** 10.1101/753442

**Authors:** Philippe Carmona, Sylvain Gandon

**Affiliations:** Laboratoire de Mathématique Jean Leray, Université de Nantes, 2 Rue de la Houssinière F-44322 Nantes Cedex; Centre D’Écologie Fonctionnelle et Évolutive, UMR 5175 C.N.R.S, 119 Route de Mende, 34293 Montpellier 5

**Keywords:** Pathogen emergence, seasonal environments, optimal control, Zika virus, epidemiology

## Abstract

Many infectious diseases exhibit seasonal dynamics driven by periodic fluctuations of the environment. Predicting the risk of pathogen emergence at different points in time is key for the development of effective public health strategies. Here we study the impact of seasonality on the probability of emergence of directly transmitted pathogens under different epidemiological scenarios. We show that when the period of the fluctuation is large relative to the duration of the infection, the probability of emergence varies dramatically with the time at which the pathogen is introduced in the host population. In particular, we identify a new effect of seasonality (the *winter is coming* effect) where the probability of emergence is vanishingly small even though pathogen transmission is high. We use this theoretical framework to compare the impact of different control strategies on the average probability of emergence. We show that, when pathogen eradication is not attainable, the optimal strategy is to act intensively in a narrow time interval. Interestingly, the optimal control strategy is not always the strategy minimizing *R*_0_, the basic reproduction ratio of the pathogen. This theoretical framework is extended to study the probability of emergence of vector borne diseases in seasonal environments and we show how it can be used to improve risk maps of Zika virus emergence.

## 1 Introduction

The development of effective control strategies against the emergence or re-emergence of pathogens requires a better understanding of the early steps leading to an outbreak Diekmann et al. [2013], Heesterbeek et al. [2015], Keeling and Rohani [2011], Woolhouse [2002]. Classical models in mathematical epidemiology predict that whether or not an epidemic emerges depends on 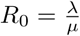 the basic reproduction ratio of the pathogen, where *λ* is the *birth rate* of the infection (a function of the transmission rate and the density of susceptible hosts) and *µ* is the *death rate* of the infection (a function of the recovery and mortality rates). In the classical deterministic description of disease transmission, the pathogen will spread if *R*_0_ > 1 and will go extinct otherwise (Figure 1). This deterministic description of pathogen invasion relies on the underlying assumption that the initial number of introduced pathogens is large. The early stages of an invasion are, however, typically characterized by a small number, *n*, of infected hosts. These populations of pathogens are thus very sensitive to demographic stochasticity and may be driven to extinction even when *R*_0_ > 1. The probability of emergence 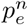 refers to the probability that, after the introduction of *n* infected hosts, a non-evolving pathogen avoids initial extinction and leads to an epidemic. Under the reasonable assumption that the initial spread of directly transmitted disease follows a one dimensional birth-death branching process the probability of emergence is zero when *R*_0_ < 1 and, when *R*_0_ > 1, it is equal to Allen and van den Driessche [2013], Allen and Jr [2012], Diekmann et al. [2013], Gandon et al. [2013], Heesterbeek et al. [2015]:

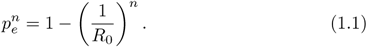

**Figure 1:**
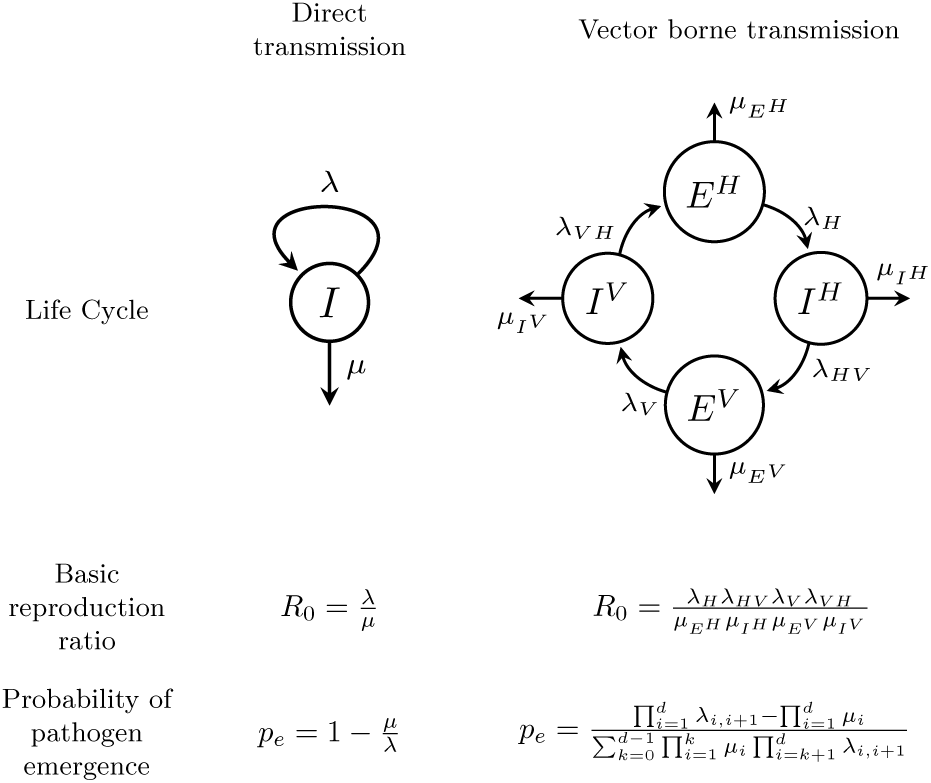
Transmission mode and pathogen emergence without seasonality. In a direct transmission model pathogen dynamics is driven by the birth rate *λ* and the death rate *µ* of a single infected compartment *I*. In a vector borne transmission model pathogen dynamics is driven by the birth rates and death rates of multiple compartments: exposed and infected humans (*E*^*H*^, *I*^*H*^), exposed and infected mosquito vectors (*E*^*V*^, *I*^*V*^). In the absence of seasonality (i.e. no temporal variation in birth and death rates) the basic reproduction ratio *R*_0_ can be expressed as a ratio between birth and death rates. The probability of emergence *p*_*e*_ after the introduction of a single infected individual can also be expressed as a function of these birth and death rates. With vector borne transmission this probability of emergence depends on which infected host is introduced (Figure S5). Here we give the probability of emergence after the introduction of a single human exposed to the pathogen, *E*^*V*^, and where the index *i* refers to the four consecutive states of the pathogen life cycle (see supplementary information).

The above results rely on the assumption that birth and death rates of the infection remain constant through time (i.e. time homogeneous branching process). Many pathogens, however, are very sensitive to fluctuations of the environment. For instance, the fluctuations of the temperature and humidity have been shown to have a huge impact on the infectivity of many viral pathogens like influenza [Shaman and Kohn, 2009] and a diversity of other infectious diseases [Altizer et al., 2006, Martinez, 2018]. In addition, many pathogens rely on the presence of arthropod vectors for transmission and the density of vectors is also very sensitive to environmental factors like temperature and humidity [Mordecai et al., 2013]. To account for these environmental variations, the birth and death rates are assumed to be functions of time: *λ*(*t*) and *µ*(*t*), respectively. The basic reproduction number is harder to compute but the probability of emergence *p*_*e*_(*t*_0_) when one infected individual is introduced (i.e. *n* = 1) at time *t*_0_ is well known (see e.g. [Kendall, 1948] or [Bailey, 1964, Chapter 7]):

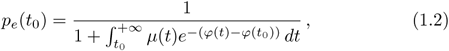

with 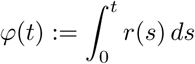 where *r*(*t*) = *λ*(*t*) − *µ*(*t*) is the malthusian growth rate of the pathogen population at time *t*. Because we are interested in seasonal variation we can focus on periodic scenarios where both *λ*_*T*_ (*t*) and *µ*_*T*_ (*t*) have the same period *T*, one year. In this case, the basic reproduction number has been computed in Bacaër and Guernaoui [2006], Diekmann et al. [2013] as the spectral radius of the next generation operator, and is the ratio of time averages of birth and death rates:

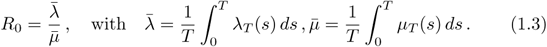

When *R*_0_ < 1 the pathogen will never produce major epidemics and will always be driven to extinction. When *R*_0_ > 1, however, a pathogen introduced at a time *t*_0_ may escape extinction. In this case the probability of emergence can also be expressed as a ratio of average birth and death rates, but with different weights:

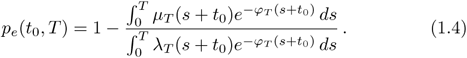

Note that this quantity refers to the probability of major epidemics, the probability that the pathogen population does not go extinct. Minor epidemics are likely to outburst if the pathogen is introduced during the high transmission season but those outbreaks do not count as major epidemics if they go extinct during the low transmission seasons.

In the following we show that very good approximations of the probability of pathogen emergence can be derived from this general expression when the period is very large (or very small) compared to the duration of the infection. These approximations give important insights on the effect of the speed and the shape of the temporal fluctuations of the environment on the probability of pathogen emergence. We use this theoretical framework to determine optimal control strategies that minimize the risk of pathogen emergence. We provide clear cut recommendations in a range of epidemiological scenarios. We also show how this theoretical framework can be extended to account for the effect of seasonality in vector borne diseases. More specifically, we use this model to estimate the probability of Zika virus emergence throughout the year at different geographic locations.

## 2 Results

### 2.1 Emergence of directly transmitted pathogens

For the sake of simplicity we start our analysis with a directly transmitted disease with a constant duration of infection, *µ*(*t*) = *µ*, but with seasonal fluctuations of the transmission rate, *λ*(*t*). This epidemiological scenario may capture the seasonality of many infectious diseases. For instance, increased contact rates among children during school terms has been shown to have a significant impact on the transmission of many childhood infections Fine and Clarkson [1982], Finkenstädt and Grenfell [2000]. Seasonal fluctuations in temperature and humidity can also drive variations in the survival rate of many viruses and result in seasonal variations in transmission rates Cook et al. [1990], Pascual et al. [2002].

Both the speed and the amplitude of the fluctuations of *λ*(*t*) can affect the probability of pathogen emergence. Yet, when the period *T* of the fluctuations is short compared to the duration 1*/µ* (e.g. fluctuations driven by diurnal cycles are fast), the probability of pathogen emergence can be approximated by (Figure 2E and 2F):

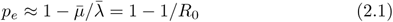

**Figure 2:**
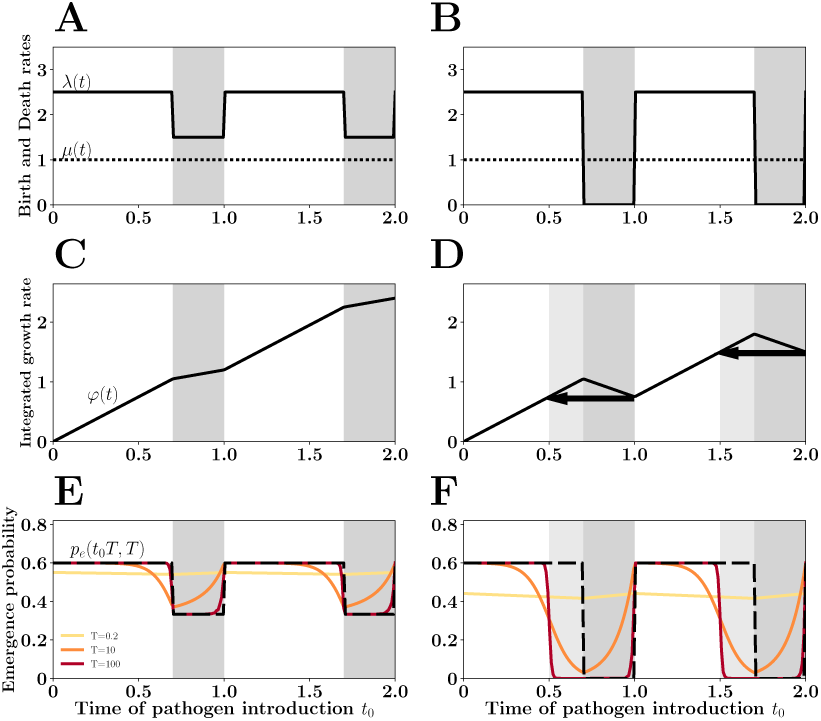
The *winter is coming* effect. Pathogen *birth rate* (i.e. transmission rate) *λ*(*t*) is assumed to vary periodically following a square wave (A and B). During a portion 1*-γ* of the year transmission is maximal and *λ*(*t*) = *λ*_0_. In the final portion of the year *λ*(*t*) drops (low transmission season in gray). Pathogen *death rate µ*(*t*) (a function of recovery and death rates of the infected host) is assumed to be constant and equal to 1 in this figure. When the net growth rate of the pathogen remains positive in the low transmission season (*λ*(*t*) > *µ*(*t*), A, C and E) the probability of emergence of a pathogen introduced at time *t*_0_ can be well approximated by 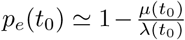 (dashed line in E and F) if the duration of the infection is short relative to the period *T* of the fluctuation (E). In contrast, if the low transmission season is more severe (*λ*(*t*) < *µ*(*t*), B, D and F), the negative growth rate *ϕ*(*t*) of the pathogen population during this period creates a demographic trap and reduces the probability of emergence at the end of the high transmision season. This *winter is coming* effect is indicated with black arrow in (D). This effect is particularly pronounced when the period of the fluctuations of the environment is large relative to the duration of the infection (i.e., when *T* is large, F).

In other words, the probability of emergence does not depend on the timing of the introduction event and it is only driven by the average transmission rate.

When the fluctuations are slower, however, the probability of pathogen emergence does depend on the timing of the introduction. The probability of emergence drops with the transmission rate (Figure 2E and 2F). When the period of the fluctuation is long, a natural approximation is:

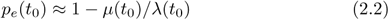

This is a very good approximation whenever the birth rate of the infection remains higher than the death rate throughout the year (i.e., *λ*(*t*) > *µ*(*t*), Figure 2). However, when *λ*(*t*) can drop below *µ*(*t*), the above approximation fails to capture the dramatic reduction of the probability of emergence occurring at the end of the high transmission season. When the introduction time of the pathogen is shortly followed by a low transmission season, the introduced pathogen is doomed because it will suffer from the bad times ahead (see supp info). We call this the *winter is coming* effect (see geometric interpretation in Figure 2D and 2F). We explore this effect in Figure S1 of supp info, under different types of seasonal variations: square waves and sinusoidal waves. As expected, the *winter is coming* effect is particularly pronounced when the period of the fluctuations are long (Figure 2F).

#### Optimal control

Our theoretical framework can be used to identify optimal control strategies. The objective is to minimize the average probability of emergence under the assumption that the introduction time is uniformly distributed over the year:

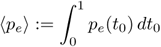

Control is assumed to act via an instantaneous reduction *ρ*(*t*) of the transmission rate of the pathogen: *λ*_*ρ*_(*t*) = *λ*(*t*)(1 − *ρ*(*t*)). We also assume that higher control intensity is costly and we define the cost of a given control strategy as a function of the intensity and the duration of the control:

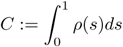

More explicitly, we assume that the control strategy is governed by three parameters: *t*_1_ and *t*_2_, the times at which the control starts and ends, respectively, and *ρ*_*M*_ the intensity of control during the interval [*t*_1_, *t*_2_]. The cost of such a control strategy is thus: *C* = (*t*_2_ *- t*_1_)*ρ*_*M*_. For a given investment in disease control *C*, what are the values of *t*_1_, *t*_2_ and *ρ*_*M*_ that minimize the average probability of emergence ⟨*p*_*e*_⟩?

We first answer this question when the fluctuation of transmission is a square wave where *λ*(*t*) oscillates between *λ*_0_ (for a fraction 1 − *γ* of the year) and 0, while *µ*(*t*) = 1 throughout the year (Figure 3). For instance, such periodicity may be driven by school terms Grassly and Fraser [2006]. The basic reproduction after control in the high transmission period is equal to *R*_0_ *- C* (see supp info). In other words, under this scenario, when the investment in control reaches a threshold (i.e. when *C* > *R*_0_ − 1) the basic reproduction (after control) of the pathogen drops below one and the probability of emergence vanishes. However, when this level of control is unattainable, the timing and the intensity of the optimal control strategy that minimizes ⟨*p*_*e*_⟩ remains to be determined. Let us first consider a naïve strategy where control is applied throughout the high transmission season. This naive strategy can be outperformed by alternative strategies where the control is applied more intensely but in a limited portion of the high transmission season (Figures 3A, 3C, 3E and S2). Indeed, this optimal strategy maximizes the *winter is coming* effect. Figure 4A shows that there is a broad range of strategies that minimize ⟨*p*_*e*_⟩ (within the dotted red curve).

**Figure 3:**
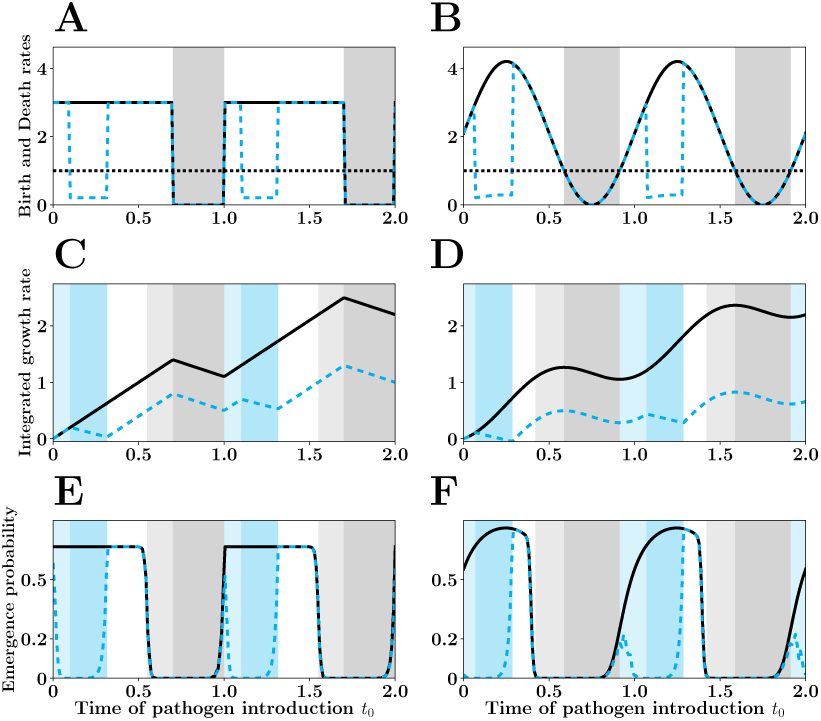
Optimal Control for square wave (A, C and E) and sinusoidal birth rates (B, D and F). In A and B we plot The pathogen birth rate before (black line) and after the optimal control (dashed blue line) which minimizes the mean emergence probability < *p*_*e*_ > (see also Figure 4). The square wave assumes that *λ*(*t*) = 3 **1**_(0<*t*<0.7)_. The sinusoidal wave assumes that *λ*(*t*) = 2(1 + sin(2*πt*)). As in Figure 2 the gray shadings refers to the low transmission season (gray) and the winter is coming effect (light gray). Similarly, we indicate the additional low transmission period induced by control (blue shading) and the additional *winter is coming* effect induced by control (light blue shading).

**Figure 4:**
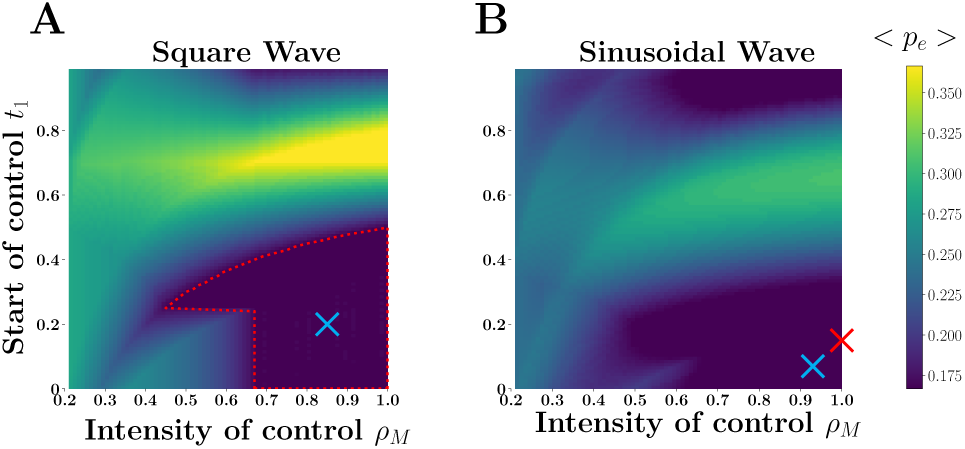
Mean probability of pathogen emergence for different control strategies with (A) square wave and (B) sinusoidal wave fluctuations. We used the same scenarios as in Figure 3 and we fix the investment in control (cost of control *C* = *ρ*_*M*_ (*t*_2_ *- t*_1_) = 0.2). We explore how the intensity of control (*ρ*_*M*_) and the timing of control (between *t*_1_ and *t*_2_) affect < *p*_*e*_ >, the mean probability of pathogen emergence (lighter shading refers to higher values of < *p*_*e*_ >). For the square wave scenario we identify a range of optimal strategies withing the dotted red curve where < *p*_*e*_ > is minimized. The optimal strategies used in Figure 3 are indicated with a blue cross for both the square wave (A) and the sinusoidal wave (B). The minimal and maximal value for < *p*_*e*_ > are: 0.166*-*0.366 (square wave) and 0.085 − 0.31 (sinusoidal wave). For the square wave (A), *R*_0_ = 1.5 does not depend on the timing and the intensity of the control. For the sinusoidal wave (B), there is a single strategy minimizing *R*_0_, namely *R*_0_ = 1.28 for *t*_1_ = 0.15 and *ρ*_*M*_ = 1.0, marked with a red cross in B. With the sinusoidal wave there is a single control strategy minimizing < *p*_*e*_ > for *t*_1_ = 0.07 and *ρ*_*M*_ = 0.93 (blue cross in B).

Second, we consider a seasonal environment where *λ*(*t*) follows a sinusoidal wave, while *µ*(*t*) = 1 throughout the year (Figures 3B, 3D and 3F). Such periodicity may arise with more gradual changes of the abiotic environment driven by climatic seasonality Grassly and Fraser [2006]. Under this scenario, pathogen transmission varies continuously and the basic reproduction after control does depend on the time at which control is applied. The basic reproduction ratio is minimized when the intensity of control is maximal (*ρ*_*M*_ = 1) in a time interval centered on the time at which pathogen transmission reaches its peak (red cross in Figure 4B). In contrast, the optimal control strategy that minimizes ⟨*p*_*e*_⟩ starts earlier, lasts longer and is a bit less intense (blue cross in Figure 4B).

### 2.2 Emergence of vector borne pathogens

Next we want to expand the above analysis to a more complex pathogen life cycle. Indeed, many emerging pathogens are vector borne Jones et al. [2008a], Woolhouse [2002]. Arboviruses, for instance, use different mosquito species as vectors and are responsible for major emerging epidemics in human populations Weaver and Barrett [2004]. In the following, we use a classical epidemiological model of Zika virus transmission which has been parameterized using empirical data sets to determine the probability of emergence under various regimes of seasonality (see sup info). In this model, the pathogen may appear in four different states (Figure 1): exposed and infectious mosquitoes (*E*^*V*^ and *I*^*V*^), exposed and infectious humans (*E*^*H*^ and *I*^*H*^). The stochastic description of this epidemiological model yields a four dimension multi-type birth-death branching process (see supp info). In the absence of seasonality (homogeneous case) the basic reproduction ratio of the pathogen is the ratio of the product of birth rates by the product of the death rates (Figure 1). The probability of emergence after the introduction of a single infected host in state *∈* (*E*^*H*^, *I*^*H*^, *E*^*V*^, *I*^*V*^):

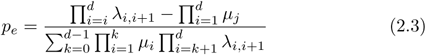

where the index *i* refers to the *d* consecutive states of the pathogens, starting with the state in which the pathogen is introduced. Note that the state of the introduced individual can have a huge impact on the probability of pathogen emergence (Figure S5).

Seasonality can drive pathogen transmission through the fluctuations of the available density of the mosquito vector. Following Zhang et al. [2017] we assume that mosquito density fluctuates with temperature and is maximal at *T*_*opt*_, the optimal temperature for mosquito reproduction (see sup info). The rate 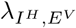 at which mosquitoes are exposed to the parasite is directly proportional to *N*_*V*_ */N*_*H*_. In such a fluctuating environment the *R*_0_ is the spectral radius of the next generation operator, see Bacaër and Guernaoui [2006], Diekmann et al. [2013] but there is no analytic expression for *R*_0_. Yet, it is tempting to use equation (2.3) with the birth and death rates functions of the introduction time *t*_0_, to obtain an approximation *p*_*e*_(*t*_0_) for large periods. The exact probability of emergence can be efficiently computed numerically thanks to the seminal work of Bacaër and Ait Dads [2014]. Figure 5 explores the difference between this naive expectation and the exact value of the probability of emergence. Crucially, we recover the same qualitative patterns observed in the direct transmission model. In particular, we notice that when the product of birth rates remains higher than the product of death rates the naive expectation for the probability of emergence is not too far from the exact value of *p*_*e*_(*t*_0_). However, when seasonality induces more pronounced drops in transmission, we recover the *winter is coming* effect where the probability of emergence can be very low before the low transmission season (Figure 5D). It is also possible to identify numerically the optimal control strategies minimizing the probability of Zika emergence (Figure S6).

**Figure 5:**
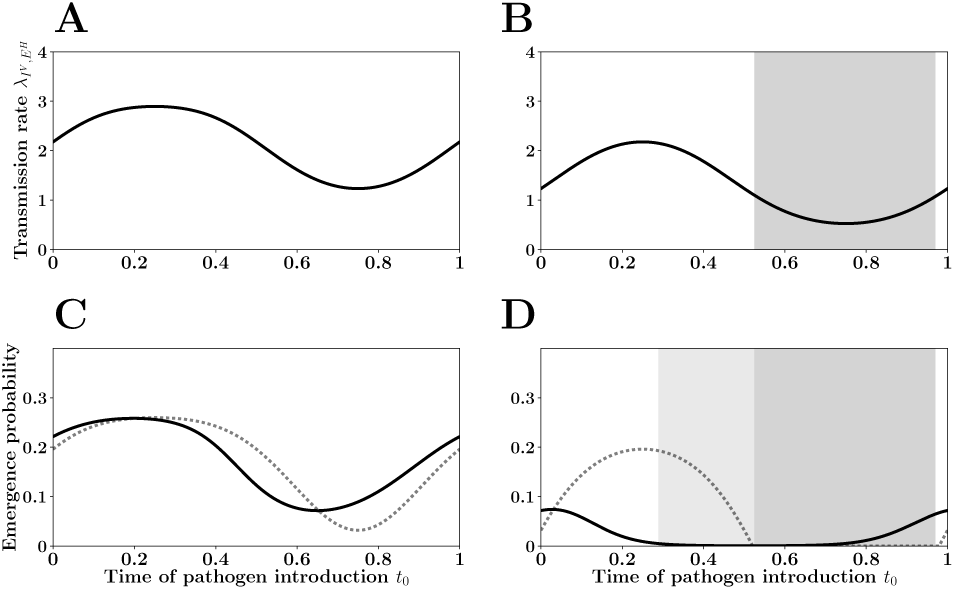
Probability of Zika emergence across space and time. The top figures (A and B) show the seasonal variations in 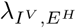, the transmission rate from humans to the vectors because of the fluctuations the density of vectors in two habitats (this illustrates the effect of *space* on Zika emergence): a minor variation in mean temperature, 29 ^°C^ (A and C) versus 27 ^°C^ (B and D), has a massive impact on transmission and, consequently, on pathogen emergence. In C and D we illustrate the effect of the *time* of introduction *t*_0_ on Zika emergence. The dotted black line refers to the naive expectation for the probability of pathogen emergence at time *t*_0_ if all the rates were constant and frozen at their *t*_0_ values (see (2.3)). The gray shading in B and D refers to the low transmission season where the product of the transmission rates is lower than the product of death rates (see supplementary information). The exact probability of emergence *p*_*e*_(*t*_0_*T, T*) is indicated as a solid black line. Higher seasonality (B and D) increases the discrepancy between the naive expectation and the exact value of the probability of pathogen emergence. This discrepancy is due to the *winter is coming* effect (light gray shading in D). See table S1 (model I) for parameter values.

## 3 Discussion

The effect of seasonality on the probability of pathogen emergence depends critically on the duration of the infection 1*/µ* relative to the period *T* of the fluctuation. When the period of the fluctuation is small (i.e., *T* < 1*/µ*) the environment changes very fast and the probability of emergence does not depend on the timing of pathogen introduction but on the average transition rates of the pathogen life cycle. When the period of the fluctuation is large (i.e., *T* > 1*/µ*) the probability of emergence varies with the timing of pathogen introduction. This probability drops when the pathogen is introduced at a point in time where conditions are unfavorable (low transmission and/or high recovery rates). More surprisingly, we show that the probability of pathogen emergence can also be very low in times where conditions are favourable if they are followed by a particularly hostile environment. This *winter is coming* effect results from the existence of adverse conditions that introduce demographic traps (where the net reproduction rate is negative) and pathogen emergence is only possible if the pathogen introduction occurs sufficiently far ahead of those traps. Note that our approach neglects the density dependence that typically occurs after some time with major epidemics. Our probability of pathogen emergence thus provides an upper approximation of the probability emergence. Indeed, with density dependence the size of the pathogen population may be too small to survive even very shallow demographic traps. In the supplementary information (section 7) we show how such density dependence can magnify the *winter is coming* effect.

Understanding this effect allows to identify the optimal deployment of control strategies minimizing the average probability of pathogen emergence in seasonal environments. We identified optimal control strategies in different epidemiological scenarios under the assumption that the introduction time is homogeneous (Figures 3, 5, S1, S4). This assumption can be readily modified to take into account temporal variations in the probability of introduction events, which yields different recommendations for the timing of control (see section 3.3 of sup info and Figure S3).

This work can be extended to explore optimal timing of other control strategies. For instance Grassly and Fraser [2006] study the optimal timing of pulse vaccination in seasonal environment and show for a range of epidemiological scenarios that a pulse vaccination applied periodically 3 months before the peak transmission rate minimizes *R*_0_. Yet, as pointed out above, the strategy minimizing *R*_0_ may not always coincide with the strategy minimizing ⟨*p*_*e*_(*t*_0_)⟩ (see Figure 4). Indeed, we found that the probability of emergence is minimized if pulse vaccination occurs a bit sooner than the time at which *R*_0_ is minimized (3.71 instead of 3 months before the peak transmission).

So far we focused on control strategies that lower pathogen transmission. Our approach can also be used to optimize control measures that do not act on the transmission rate but on the duration of the infection. For instance, what is the optimal timing of a synchronized effort to use antibiotics to minimize bacterial pathogens emergence? We found that the timing of these treatment days have no impact on *R*_0_ but pathogen emergence is minimized when treatment occurs 1.3 months before the peak of the transmission season. This strategy creates deeper traps and results in a stronger *winter is coming* effect. Interestingly, Lee et al. [2005] explored the optimal timing of mass antibiotic treatment to eliminate the ocular chlamydia that cause blinding trachoma. Numerical simulations showed that the speed of eradication is maximized (the time to extinction is minimized) when treatment is applied 3 months before the low transmission season. A similar result was obtained by Gao et al. [2014] showing that it is best to treat against malaria in the low transmission season. The apparent discrepancy between these recommendations is driven by the use of different objective functions (pathogen emergence, speed of eradication or cumulative number of cases).

The above examples show that our analysis has very practical implications on the understanding and the control of emerging infectious diseases in seasonal environments. This theoretical framework could be used to produce maps with a very relevant measure of epidemic risk: the probability of pathogen emergence across space and time (Figure 5). Currently available risk maps are often based on integrated indices of suitability of pathogens or vectors Jones et al. [2008b], Kitron [2000], Kraemer et al. [2016], Nsoesie et al. [2016]. These quantities may be biologically relevant but the link between these quantities and the probability of pathogen emergence is not very clear. We contend that using risk maps based on *p*_*e*_(*t*_0_) would be unambiguous and more informative. More generally, the same approach could also be used to improve the prevention against invasions by nonindigenous species Lodge et al. [2006].

Experimental test of theoretical predictions on pathogen emergence are very scarce because the stochastic nature of the prediction requires massive replicate numbers. Some microbial systems, however, offer many opportunities to study pathogen emergence in controlled and massively replicated laboratory experiments Chabas et al. [2018]. It would be interesting to use these microbial systems to study the impact of periodic oscillations of the environment to mimic the influence of seasonality. Another way to explore this question experimentally would be to use data on experimental inoculation of hosts. Indeed, the experimental inoculation of a few bacteria in a vertebrate host (which could be viewed as “population” of susceptible cells) is equivalent to the introduction of a few pathogens in a host population. The outcome of these inoculations are stochastic and the probability of a successful infection (host death) is equivalent to a probability of emergence. Interestingly, some daily periodicity to bacterial infections has been found in mice Feigin et al. [1969], Shackelford and Feigin [1973]. Mice inoculated early in the morning (4am) have a higher probability of survival than mice inoculated at any other time. This pattern is likely to result from a circadian control of the vertebrate immune system Scheiermann et al. [2013] which are likely to impact the birth and death rates of bacteria. Given that the generation time of a bacteria is smaller than a day, it is not surprising to see a probability of emergence depending on the inoculation time (see equation 2.2). In other words, our work may also be used to shed some light on the stochastic within-host dynamics of pathogen infections. One could envision that simple changes in therapeutic practices that take into account the time of day may affect clinical care and could limit the risk of nosocomial infections. Our work provides a theoretical toolbox that can integrate detailed description of the periodic nature of pathogen life cycles at different spatial and temporal scales (within and between hosts, over the period of one day or one year) to time optimal control strategies.

## Acknowledgments

We thank Mike Boots and Sébastien Lion for comments on an earlier draft. Philippe Carmona acknowledges funding from C.N.R.S which supported a 50% délégation in 2019 to the CEFE in Montpellier, as a visiting professor. He also thanks the Centre Henri Lebesgue ANR-11-LABX-0020-01 for creating an attractive mathematical environment. Sylvain Gandon acknowledges support from CNRS.

## Supplementary Material

### 1 Pathogen emergence with periodic birth and death rates

For a birth and death process with constant birth and death rates (*λ* > 0, *µ* > 0), the basic reproduction number is 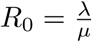 and the probability of extinction, starting initially with one individual, is 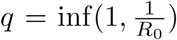. When the birth and death rate are assumed to be functions of time, noted *λ*(*t*) and *µ*(*t*), respectively, the basic reproduction number is harder to compute but the extinction probability is well known (see e.g. Kendall [1948] or [Bailey, 1964, Chapter 7]). This yields the following expression for *p*_*e*_(*t*_0_), the probability of pathogen emergence when a single infected host is introduced in the host population at time *t*_0_:

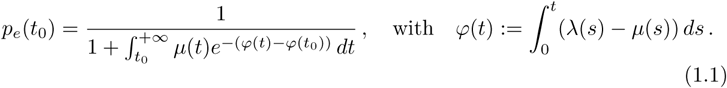

Let us now consider rates with period *T* > 0, denoted by *λ*_*T*_ and *µ*_*T*_. Accordingly, we denote 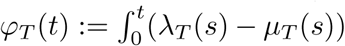 *ds* and *p*_*e*_(*t*_0_, *T*) the corresponding emergence probability.

The basic reproduction number has been derived in Bacaër and Guernaoui [2006], Diekmann et al. [2013] as the spectral radius of the next generation operator, and is the ratio of time averaged birth and death rates

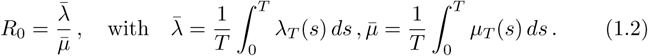

Since 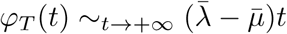, we find that *p*_*e*_(*t*_0_, *T*) = 0 if *R*_0_ ≤ 1.

If *R*_0_ > 1, we can rearrange formula(1.1) and express *p*_*e*_(*t*_0_, *T*) as a ratio of average birth and death rates, but with a weight that takes into account the average growth rate of the pathogen population:

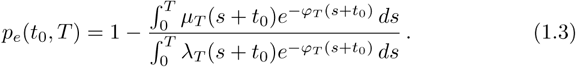

Indeed, first observe that since 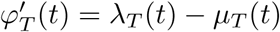 we have

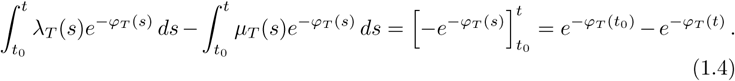

Since *φ*_*T*_ (*t*) → +∞ this implies

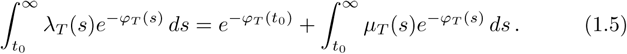

We now use periodicity to obtain, first that for integer *k*,

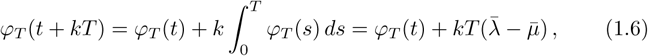

and thus

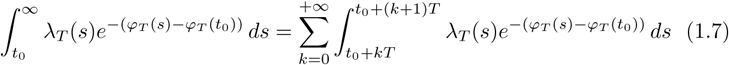

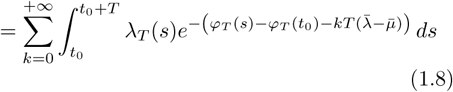

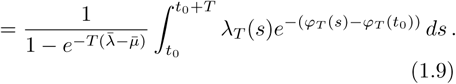

Similarly,

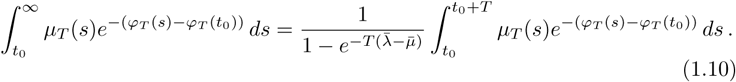

Hence,

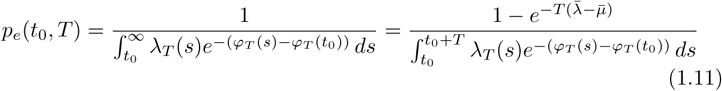

and

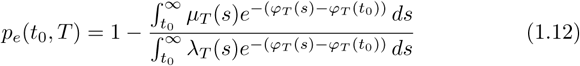

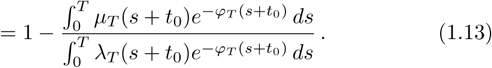

### 2 Asymptotic results for small and large periods

Under the assumption that 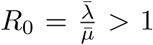 we know that *p*_e_ (*t*_0_, *T*) > 0 for all *t*_0_. In the following we rescale time so that the *T* periodic functions *λ*_*T*_, *µ*_*T*_ become 1 periodic functions defined by

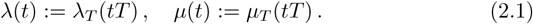

And similarly,

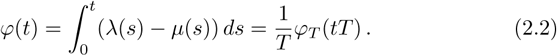

Hence, by a change of variables we obtain

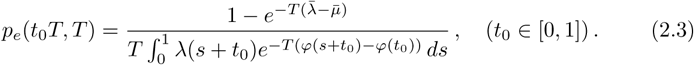

In the following we derive simpler expressions for *p*_*e*_(*t*_0_*T, T*) in the limit cases where *T* is very small or very large.

#### 2.1 Asymptotics for small periods: when *T* → 0

We see from equation (2.3) that when 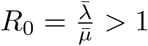 that we have

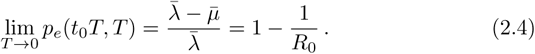

In other words when *T* → 0, we can replace the varying rates by their means. Indeed we have on one hand, as *T* → 0,

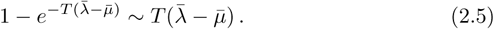

On the other hand, since *λ* has period 1,

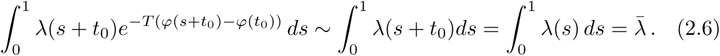

#### 2.2 Asymptotics for large periods: when *T* → +∞

We observe on various examples that for large *T*, *p*_*e*_(*t*_0_*T, T*) can sometimes be very small on subintervals of [0, *T*].

We are going to give a mathematical formulation to this observation. Define

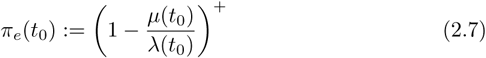

to be the *guess* we make for large periods by substituting in the formula giving the emergence probability for constant rate *λ*(*t*_0_) and *µ*(*t*_0_) to *λ* and *µ*.

##### Proposition 2.1.

*Under mild regularity assumptions on the rates λ,µ, there exists a finite set F such that for t*_0_ ∉ *F the following limit exists p*_*e*,∞_(*t*_0_):= lim_*T* →+∞_ *p*_*e*_(*t*_0_*T, T*) *and we have*

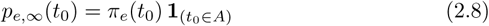

*with*

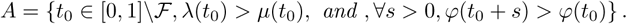

The set *A* has the following geometric interpretation. the point *t*_0_ ∈ [0, 1] is in *A* if, when the set *F* is empty,

- the rate function *ϕ* increases at *t*_0_. Indeed *ϕ*′(*t*_0_) = *λ*(*t*_0_) *- µ*(*t*_0_) > 0.
- If you look at the future starting from *t*_0_, that is you look at the function *s* → *ϕ*(*t*_0_ + *s*), you do not see a trap, that is a point lower than your starting point, that is a *s* > 0 such that *ϕ*(*t*_0_ + *s*) ≤ *ϕ*(*t*_0_).

Another way to describe this set is the following. If *t*_0_ is a local minimum of *φ*, then the points left of *t*_0_, *s* ≤ *t*_0_ where *φ* is above this local minimum, that is *φ*(*s*) ≥ *φ*(*t*_0_) are in *A*^*C*^. If you do this with all the local minimums, you get the whole of *A*^*C*^.

##### 2.2.1 Proof of Proposition 2.1

The regularity assumptions on the rate functions are the following. the functions *λ* and *µ* are assumed to be piecewise *C*^1^, that is there exists a finite set of points 0 = *x*_1_ < *x*_2_ < … < *x*_*n*_ = 1 in [0, 1] such that *λ,µ* are *C*^1^ on the segments (*x*_*i*_, *x*_*i*+1_) and *λ, λ*′, *µ, µ*′ have left and right limits at the *x*_*i*_’s. Furthermore, we assume that for every *i*, the set {*t*: *φ*(*t*) = *φ*(*x*_*i*_)} is finite, with *φ*(*t*) = ∫0 *t*(*λ*(*s*) − *µ*(*s*)) *ds*. Then, Proposition 2.1 is an immediate consequence of the two well known results on Laplace integrals, which we recall here (see e.g. Faraut [2006], section VII.3)

Let *f* be a locally bounded measurable function, and *φ* a non negative measurable function, both defined on [*a, b*]. For *λ* > 0 we let

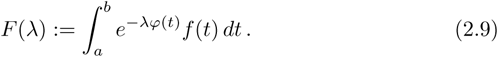

If, when *t* ↓ *a*, we have *f* (*t*) = *l* + *o*(1) and *φ*(*t*) = *α* + (*t − a*)*β* + *o*(*t*) with *β* > 0, then

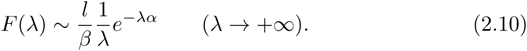

(this result is thus true if *f* is right continous at *a* and *φ* has a positive right derivative at *a*).

If when *t* ↓ *a*, we have *f* (*t*) = *l* + *o*(1) and 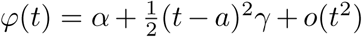 with *γ* > 0, then

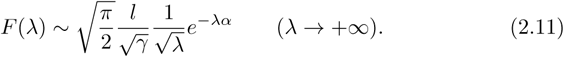

### 3 Case studies

#### 3.1 Square wave

We assume that death rate is constant *µ*(*t*) = 1 and *λ*(*t*) = *λ*_0_ **1**_(0<*t*<1*-γ*)_ with *λ*_0_ > 1. Assume furthermore that *R*_0_ > 1 that is 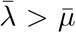 with 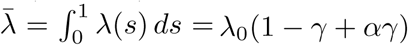 and 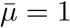.

The limiting probability of emergence is

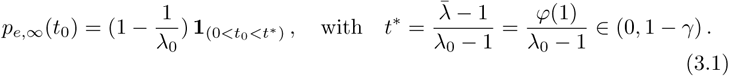

Indeed, the function *φ* is the 1 periodic function corresponding to the hat function of Figure S1, and *t*^*^ is the only solution in (0, 1) of *φ*(*t*_0_) = *φ*(1): for *t*_0_ *∈* (*t*^*^, 1) there exists *s* > 0 such that *φ*(*s* + *t*_0_) < *φ*(*t*_0_) and for *t ∈* (0, *t*^*^), for any *s* > 0, *φ*(*s* + *t*_0_) > *φ*(*t*_0_).

**Figure S1:**
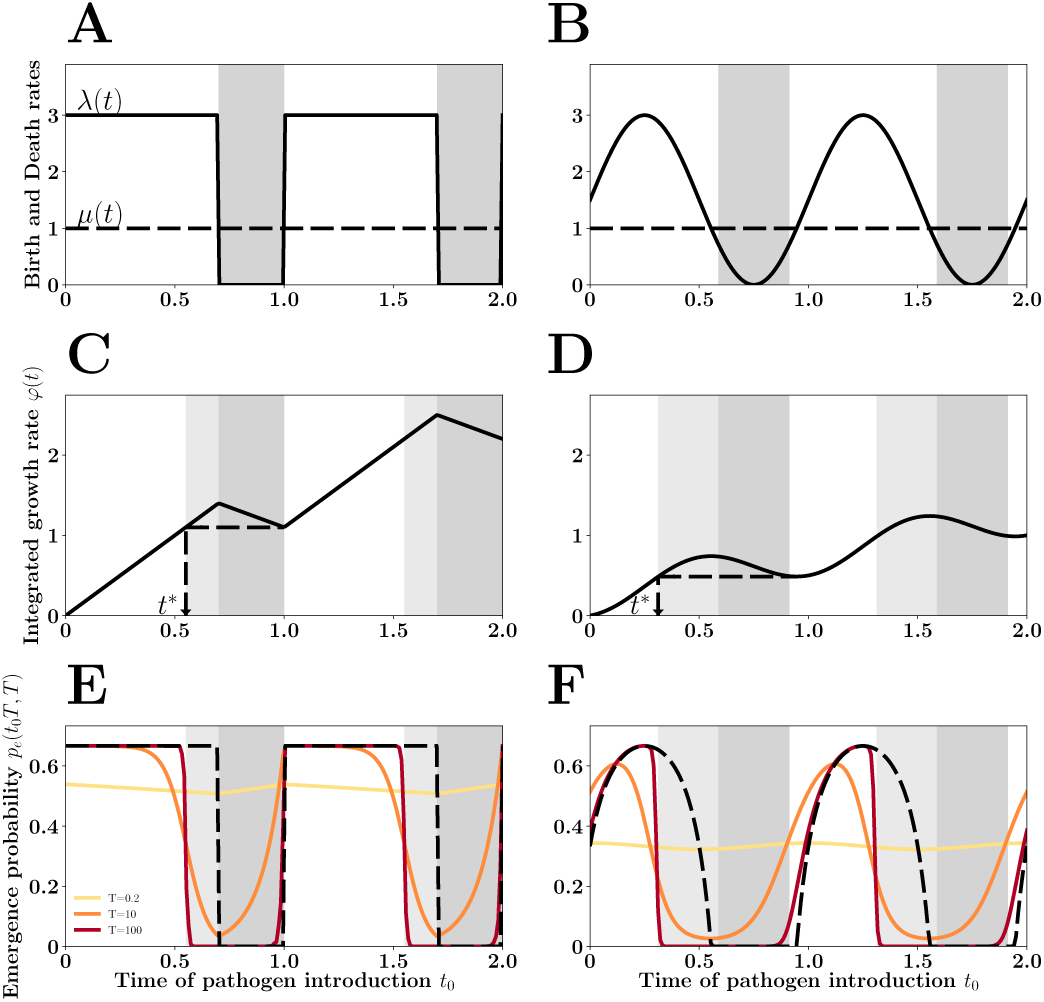
The *winter is coming* effect. Pathogen *birth rate λ*(*t*) (i.e. transmission rate) is assumed to vary periodically following a square wave (figures A, C and E) or a sinusoidal function (figures B, D and F). Pathogen *death rate µ*(*t*) (a function of recovery and death rates of the infected host) is assumed to be constant and equal to 1 in this figure. In the low transmission season we assume that transmission can become lower than the death rate (*λ*(*t*) < *µ*(*t*)). The negative growth rate of the pathogen population during this period (*r*(*t*) < 0) creates a demographic trap (a drop in *φ*(*t*), figures C and D) and reduces the probability of emergence at the end of the high transmision season (figures E and F). This *winter is coming* effect is indicated with a light gray shading between time *t*^*^ and the start of the low transmission season (figures C, D, E and F). This effect is particularly pronounced when the period of the fluctuations of the environment is large relative to the duration of the infection (i.e., when *T* is large, figures E and F).

##### 3.1.1 Control strategies

We consider control strategies that lower the birth rate *λ*(*t*) by a factor *ρ*(*t*).

More precisely we replace *λ*(*t*) by

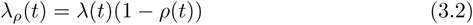

with *ρ*(*t*) *∈* [0, 1]. Accordingly *φ*(*t*) is replaced by 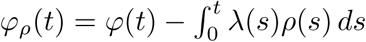 and the probability of emergence *p*_*e*,∞,*ρ*_(*t*_0_) is decreased. We measure the quality of the control strategy by averaging this quantity

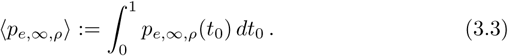

Indeed ⟨*p*_*e*,∞,*ρ*_⟩ is the mean probability of emergence for an infected who arrives uniformly in the period.

The *cost* of the control strategy is

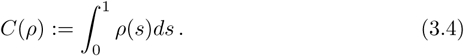

We are looking for strategies with a given cost *C* = *C*(*ρ*) which minimize ⟨*p*_*e*,∞,*ρ*_ ⟩. We shall assume from now on thath the control strategies are of the type

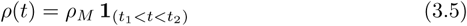

with 0 ≤ *ρ*_*M*_ ≤ 1 and 0 ≤ *t*_1_ < *t*_2_ < 1. It should be obvious that the optimal strategies should satisfy [*t*_1_, *t*_2_] ⊂ [0, 1 ≢ *γ*], and we shall assume this is the case from now on. Indeed, lowering *λ*(*t*) when it is already 0 is useless.

Hence, the basic reproduction number of these control strategies is *constant* and is

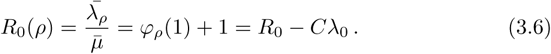

Of course, if 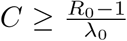, then *R*_0_(*ρ*) ≤ 1, there is almost sure extinction for all these strategies and we are done. We shall therefore assume that 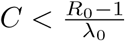 and this implies in particular that *C* ≤ 1 − *γ*.

Then, for a fixed large period *T*, one such strategy shall be close to optimal, since the extinction probabilities converge very fast.

###### Proposition 3.1.

*Among the control strategies of type* (3.5) *with* [*t*_1_, *t*_2_] ⊂ [0, 1 − *γ*] *and fixed cost* 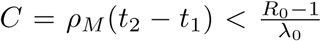, *there exists a continuum of optimal strategies that have all the minimal mean emergence probability*

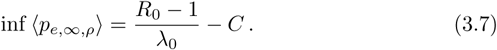

The proof is straightforward, but tedious, and is done in section 8. In the course of the proof one can see that any strategy with cost *C* such that

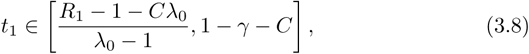

and *t*_2_≤1*- γ* is optimal.

It is natural to compare these optimal strategies to the *naive strategy* that lowers *λ* on the whole interval [0, 1 *- γ*], that is *t*_1_ = 0, *t*_2_ = 1 *- γ*:

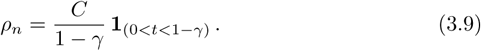

We have

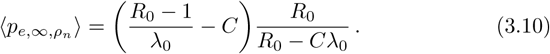

The comparison of the 〈*p*_*e*_ラ obtained is done in figure S2 where we see the curves corresponding to (3.7) and (3.10). In contrast to the *naive strategy* the optimal control strategy allows to decrease the mean probability of emergence with higher investment in control.

**Figure S2:**
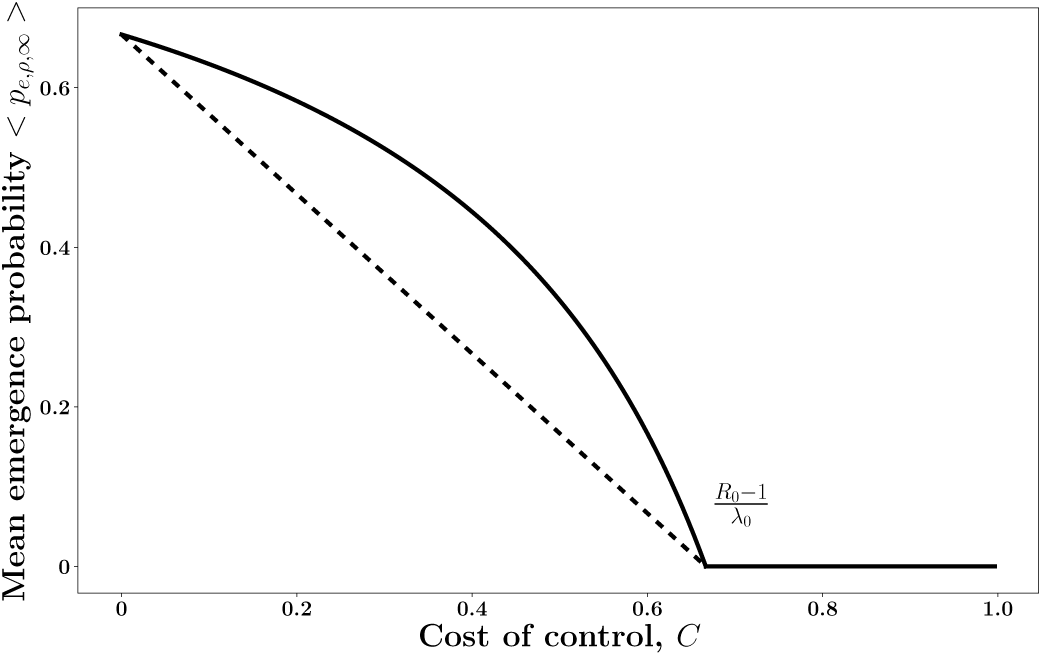
The mean probability of emergence drops with higher investment in control. The efficacy of control varies wih the timing of the intervention. The *naive strategy* (in solid black, equation (3.10)) versus the *optimal strategy* (in dashed black, equation (3.7)) when the fluctuations follow a square wave (see also figure 4 in the main text). Parameters: *λ*_0_ = 3.0, *µ* = 1, *γ* = 0.3, *R*_0_ = 2.1

### 3.2 Sinusoidal wave

We assume that death rate is constant *µ*(*t*) = 1 and that the birth rate is sinusoidal

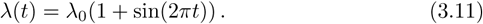

We can compute numerically the emergence probability *t*_0_ − ⟨*p*_*e*_(*t*_0_*T, T*) ⟩ and its mean. We observe that the *winter is coming* effect still applies, see figure S1

The *control strategies* are of the type

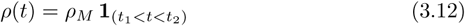

with *ρ*_*M*_ *∈* [0, 1] and 0 ≤*t*_1_ ≤*t*_2_ ≤2. We are going to compare control strategies with the same fixed cost *C* > 0. We have

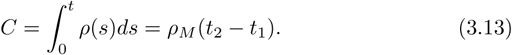

Since Δ*t* = *t*_2_ *- t*_1_ ≤ 1, we shall assume that *C* ≤ *ρ*_*M*_. We decided to parameterize the control strategies by (*t*_1_, *ρ*_*M*_) *∈* [0, 1] *×* [*C*, 1].

In contrast with the square wave fluctuation scenario, the basic reproduction number *R*_0_(*ρ*) varies with the time at which control is applied during the high transmission season. Moreover, the optimal strategy, the one that has the lowest *R*_0_ is *different* from the optimal strategy that has the lowest mean emergence (Figure 4).

### 3.3 When the time of introduction is not uniformly distributed

In the above sections we assumed that the time at which the initial infected host is introduced follows a uniform distribution. In other words we minimize:

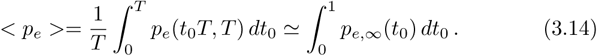

In this section we extend this approach and we allow the timing of introduction to follow a different distributiont δ(*t*). For instance we can assume that δ(*t*) is proportional to the birth rate:

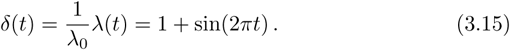

In this scenario we have to minimize:

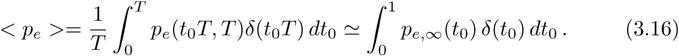

Figure S3 illustrates the influence of the distribution of the timing of pathogen introduction on mean pathogen emergence. Note that the higher rate of introduction during the high transmission season increases dramatically the risk of pathogen emergence at that time. This explains why the timing of the optimal control is slightly delayed (closer to the maximal rate of pathogen transmission/introduction). In both cases the optimal strategy uses the *winter is coming effect* to reduce mean pathogen emergence.

**Figure S3:**
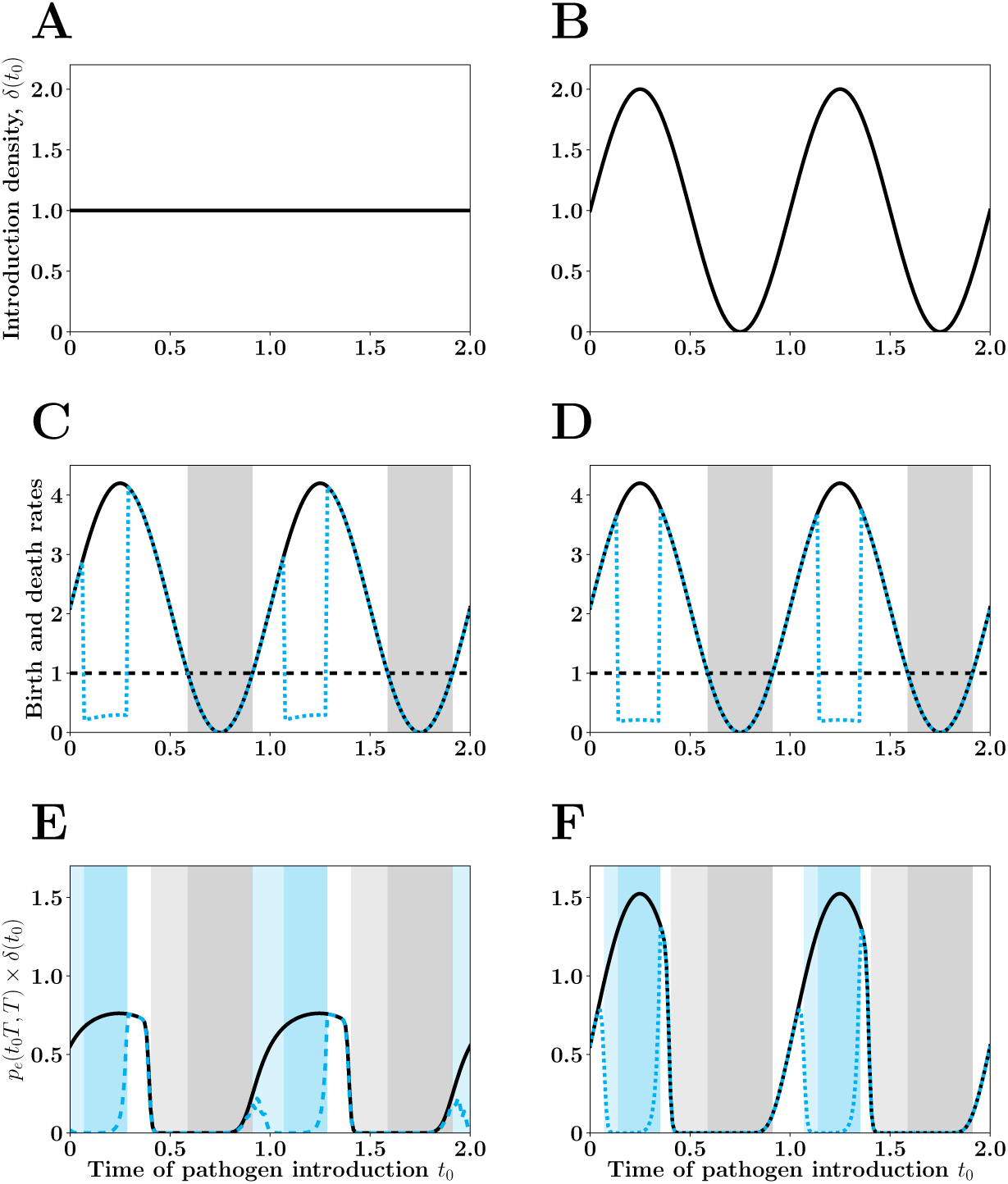
Timing of introduction and pathogen emergence. We contrast two scenarios where the time of introduction follows a homogeneous distribution (A, C and E) or a sinusoid function (B, D and F) when transmission follows a sinusoid wave. In C and D we plot the sinusoidal fluctuations of pathogen transmission before (solid black line) and after (dashed blue line) the optimal control. In E and F we plot the probability of emergence before (solid black line) and after (dashed blue line) the optimal control. The blue shading indicates the timing at which optimal control is applied and the light blue shading refers to the *winter is coming* effect induced by the control. The function plotted in the last box is not the emergence probability but the product with the density *p*_*e*_(*t*_0_*T, T*) * δ(*t*_0_) (our optimal control strategy minimizes the integral of this function).

### 4 Pathogen emergence for time varying multitype birth and death processes

Next we generalise the results obtained above in scenarios where the infected hosts may appear in different states (e.g. different host species, different states of the host). We consider a multitype birth and death processes where an individual *i* dies with rate *µ*_*i*_ and gives birth to an individual of type *j* at rate *λ*_*ij*_. These rates are assumed to be non negative but time varying, so we obtain a time inhomogeneous Markov process on ℕ^*d*^ with generator, for bounded functions *f*,

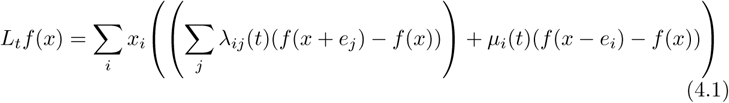

with *x ∈* ℤ^*d*^, and *e*_*i*_ the *i*-th base vector.

#### 4.1 An ODE satisfied by the extinction probabilities

Let 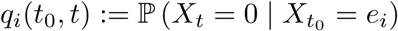 be the extinction probability at time *t* when the process starts with one individual of type *i* at time *t*_0_. Since 0 is absorbing, we know that as 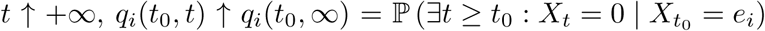, the *extinction probabilities*. The *emergence probabilities* are defined as

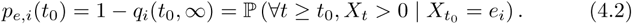

##### Proposition 4.1.

*The emergence probabilities are solutions of the system of ODE’s*

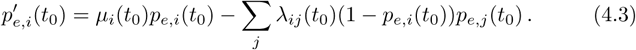

*Proof*. Assume first that *t*_0_ = 0. The first jump time, starting from *e*_*i*_, has distribution

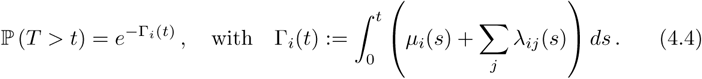

Therefore, conditioning by the value of *T*_*i*_, we get

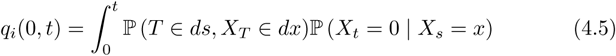

Thanks to the branching property,

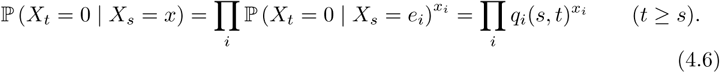

We know that conditionally on *T* = *s*, we have one offspring of type *j* with probability 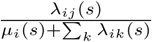 and no offspring (i.e. death of type *i*) with probability 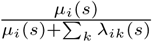 Therefore,

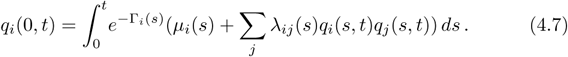

Since 0 is an absorbing set, we know that *q*_*i*_(*t*) increases to *q*_*i*_(∞), and is non negative bounded since it is a probability. By dominated convergence, we can let *t* → +∞ in the above equality to get

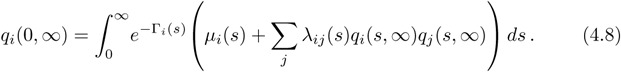

In order to obtain *q*_*i*_(*t*_0_, ∞) we replace in the preceding formula, λ_*ij*_(*s*) and *µ*_*i*_(*s*) by λ_*ij*_(*s* + *t*_0_) and *µ*_*j*_(*s* + *t*_0_).

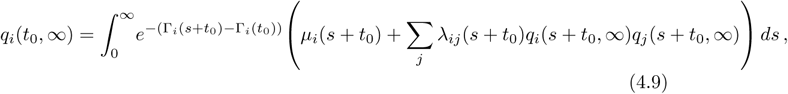

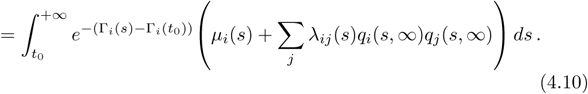

We can differentiate this equation to get

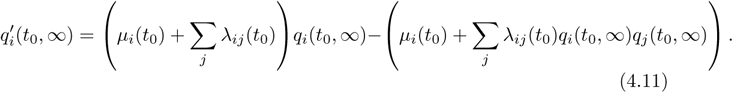

We easily check that the emergence probabilities *p*_*e,i*_(*t*_0_) = 1 −*q*_*i*_(*t*_0_, ∞) satisfy (4.3).□

##### Remark 4.2

*As usual the null function p*_*e,i*_(*t*_0_) = 0 *is always a solution of* (4.3) *and we can prove easily, by monotonicity arguments, that p*_*e,i*_(*t*_0_) *is the largest solution, in* [0, 1] *of equation* (4.3).

*Let R*_0_ *be the basic reproduction number for this birth and death process: we know, for example from Bacaër and Guernaoui [2006] that R*_0_ *>* 1 *iff ∀i, ∀t*_0_, *p*_*e,i*_(*t*_0_) *>* 0.

*Eventually, observe that the ODE* (4.3) *is a time varying Lotka-Volterra systems of equations, and we know that even in dimension* 2, *no exact solution is known*.

#### 4.2 A special case: when all rates are constant

In this section we use the general expressions derived above in the special case where all the birth and death rates are constant. We obtain the system:

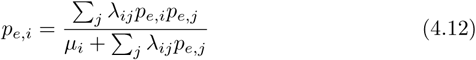

We know that when *R*_0_ *>* 1, the *p*_*e,i*_ are positive solutions of this system.

##### In dimension 1

*R*_0_ *>* 1 means λ *> µ* and 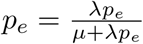 yields *p*_*e*_ = 1 − *µ/*λ.

##### In dimension *d* = 2

If *λ*_11_ = *λ*_22_ = 0 we get

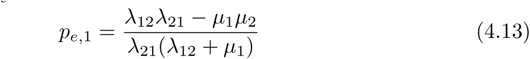

and of course we have *R*_0_ *>* 1 iff *λ*_12_*λ*_21_ − *µ*_1_*µ*_2_ *>* 0.

##### The *cyclic* case

In a *cyclic* case an individual of type *i* can give rise to a single type *j* (noted *i* + 1), and no other type can give rise to *i* + 1. In this scenario we have:

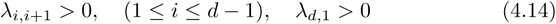

and the other *λ*_*ij*_ are 0. To simplify notations we let *λ*_*d,d*+1_ = *λ*_*d*,1_.

First observe that *R*_0_ *>* 1 is equivalent to Π *λ*_*i,i*+1_ *>* Π *µ*_*i*_.

We have the formula,

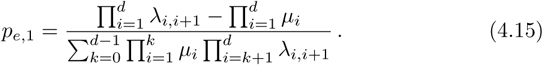

The other *p*_*e,i*_ are given by a cyclic permutation of the preceding formula. Indeed, the equation (4.12) is now, with the convention that *p*_*e,i*_ = *p*_*e,i* *mod*(*d*)_

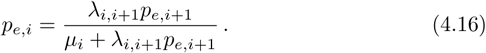

We start from *p*_*e*,1_, we then plug in the formula for *p*_*e*,2_, … and when we come back to *p*_*e*,1_ we simplify by *p*_*e*,1_ *>* 0 to get formula (4.15).

#### 4.3 The numerical approach of Bacaër and Ait Dads

In principle, there should be no problem to numerically approximate *p*_*e,i*_ a solution of (4.3). However, we do not know how to fix boundary counditions. If we knew a way to fix them, that would mean that the values of *p*_*e,i*_(0) are already known.

Bacaër and Ait Dads [2014] established an approximation of the extinction probabilities *p*_*e,i*_ by combining the method of characteristics and Kolmogorov’s formard equation. We used these to perform our computer simulations. The method is the following.

Let *τ* be a large number, at least large with respect to the period *T*. Let *Y* ^(*τ*)^ be the solution, with initial condition 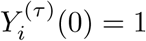, of

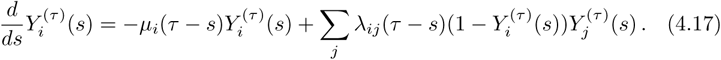

Then, when *τ* is large enough, 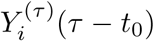 is close to *p*_*e,i*_(*t*_0_) as illustrated in figure S4: if you *reverse time* you look at the top curve from right to left that is you look at 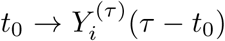 (and then you get a very close approximation to *t*_0_ → *p*_*e,i*_(*t*_0_)).

**Figure S4:**
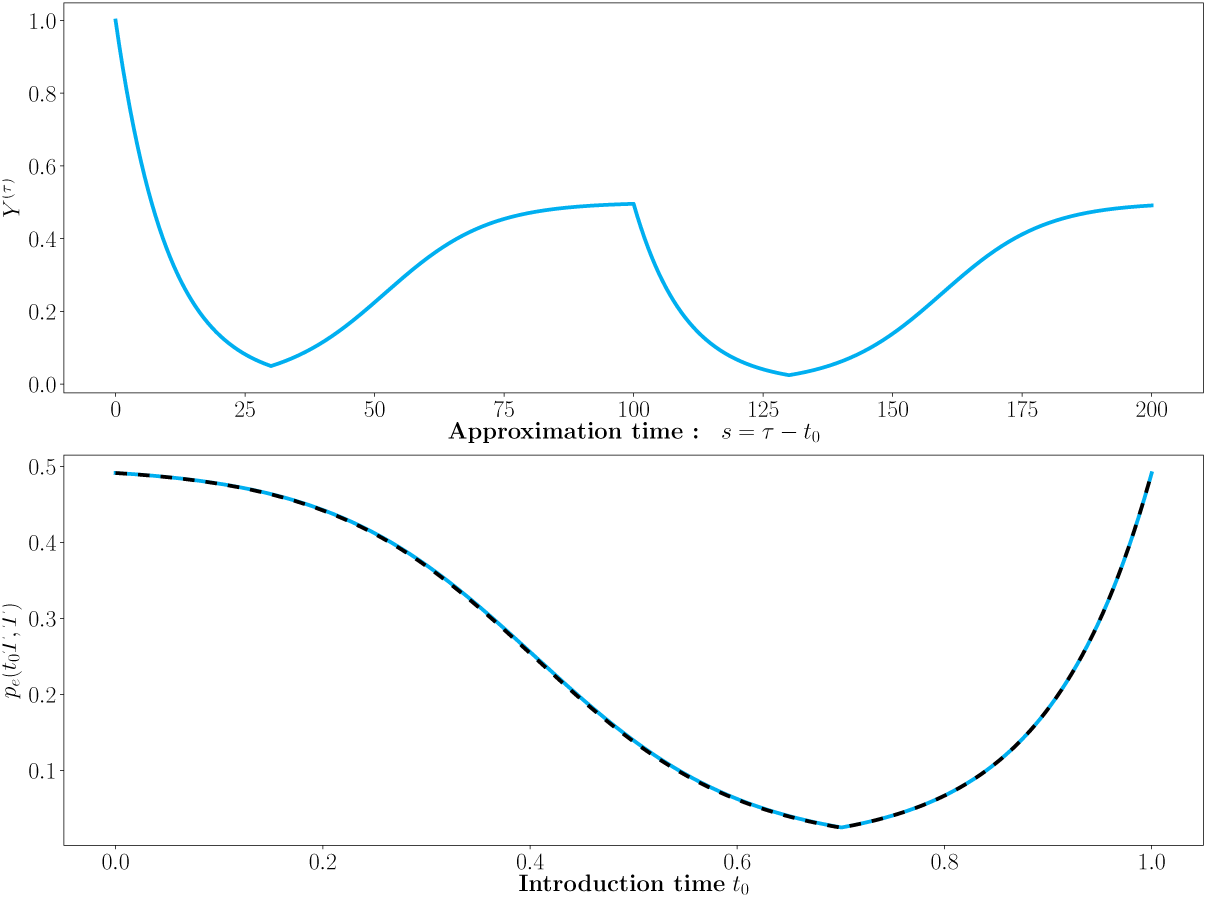
Numerical calculation of *p*_*e*_(*t*_0_*T, T*). The top graphic is *Y* ^*τ*^ (*s*) for a one dimensional birth and death process with birth rate *λ*(*s*) = 0.2 **1**_(0*<s<*0.7)_, *µ*(*s*) = 0.1, the period *T* = 100 and *τ* = 2*T*. We see that it stabilizes very fast to the periodic solution. The bottom graphic shows that the emergence probability *p*_*e*_(*t*_0_*T, T*) as a function of *t*_0_, in dotted black, coincides numerically with its approximation *Y* ^*τ*^ (*τ* − *t*_0_), in solid blue, that is the top curve taken right to left on one period.

We are now going to give a probabilistic proof of the approximation

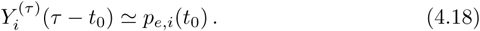

One interesting byproduct of this proof, that makes it a complement to the proof of Bacaër and Ait Dads [2014], is that it gives convergence rates.

Indeed, let us look again at extinction probabilities. As in the proof of Proposition 4.1, for 0 *≤ t*_0_ *≤ τ* we have

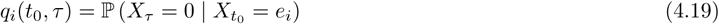

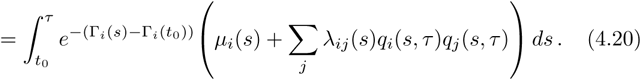

Differentiating with respect to *t*_0_, we get that *t*_0_ → *q*_*i*_(*t*_0_, *τ*) satisfy the same differential equations (4.11) as *q*_*i*_(*t*_0_, ∞), but with the boundary conditions *q*_*i*_(*τ, τ*) = 0.

In terms of emergence probabilities that means that 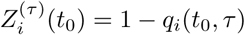 satisfy the differential equations (4.3) with the boundary condition 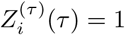. By uniqueness of ODE solutions with locally Lipschitz coefficients, this implies that

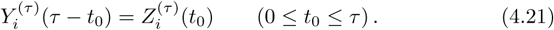

We have already observed that since 0 is an absorbing point for the branching process, *q*_*i*_(*t*_0_, ∞) = lim_*τ*→+∞_ *q*_*i*_(*t*_0_, *τ*). And this translates immediately to

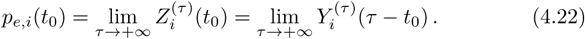

We can even show that this convergence happens exponentially fast. Indeed, let

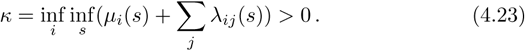

Then, remembering that 0 *≤ q*(*s, τ*) *≤* 1, we have

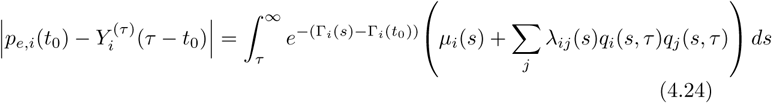

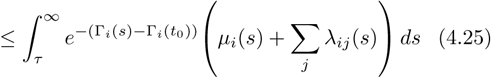

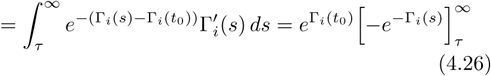

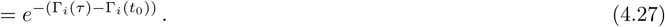

Since we have the lower bound

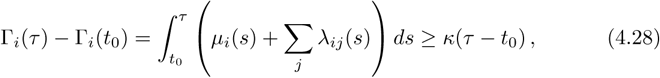

we obtain

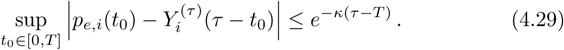

If we consider the example of figure S4, we can take *κ* = 0.1, we have *T* = 100 and we want to take *τ* = *nT*, a finite number of periods, large enough to ensure a precision of at least 10^−3^. It is enough to have *κ*(*τ* − *T*) *>* 3 log(10) and thus 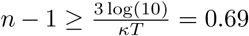. Therefore, having the approximation *Y* ^(*τ*)^ run for *τ* = 2*T* (2 periods) is largely enough.

#### 4.4 Asymptotic results for large periods: *T* → +∞

When we rescaled functions *λ*_*ij*_(*t*):= *λ*_*ij,T*_ (*tT*) and *µ*_*i*_(*t*):= *µ*_*i,T*_ (*tT*) we observe the same phenomenom as in the one dimensional scenario. The emergence probabilities *p*_*e,i*_(*t*_0_*T, T*) are very close to 0 on sub intervals. Besides, when they are not 0 they are very close to the *guess* π_*e,i*_(*t*_0_) which are obtained by substituting in the formulas giving the emergence probabilities for constant rates, the rates by their values at time *t*_0_, the time of introduction of the infected individual of type *i* (see figure 1). For example, in dimension *d* = 2, with*λ*_11_ = *λ*_22_ = 0, we have

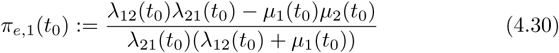

The π_*e,i*_(*t*_0_) are solutions of (4.12), where the rates *λ*_*ij*_, *µ*_*i*_ are replaced by their values at *t*_0_.

Unfortunately, we are unable to determine a set *A* such that

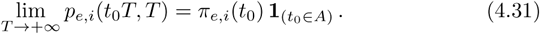

Indeed, we are unable to prove that the sequence of functions *T* → (*p*_*e,i*_(.*T, T*)) is compact in the set of continous functions from [0, 1] to R by applying Arzela-Ascoli’s theorem: the functions *p*_*e,i*_(.*T, T*) are uniformly bounded together with the functions functions *λ*_*i,j*_, *µ*_*i*_ are bounded (periodic and locally bounded), but the derivatives have a *T* factor that prevents them to be bounded. This is exactly what happens in dimension 1 where we proved that the limit was discontinuous.

However, let us consider the limit of a subsequence, i.e. assume that for a sequence *T*_*n*_ → +∞, the functions *p*_*e,i*_(*t*_0_*T*_*n*_, *T*_*n*_) converge almost everywhere on [0, 1] to the functions *α*_*i*_(*t*_0_).

##### Proposition 4.3.

*For every t*_0_ ∈ [0, 1] *such that the α*_*i*_ *are right continuous at t*_0_, *either α*_*i*_(*t*_0_) = π_*e,i*_(*t*_0_), *either α*_*i*_(*t*_0_) = 0. *In particular, whenever* π_*e,i*_(*t*_0_) = 0, *we have α*_*i*_(*t*_0_) = 0.

*Proof*. We are going to prove that *α*_*i*_(0) satisfy the same equations as the guess π_*e,i*_(0). The general result can then be deduced by considering the shifted rates *λ*_*ij*_(.) = *λ*_*ij*_(. + *t*_0_).

Let us rewrite equation (4.9), taking into account periodicity, of all the functions *λ*_*ij*_, *µ*_*i*_, *y*_*i,T*_ and so Γ_*i,T*_ (*s* + *kT*) = *k*Γ_*i,T*_ (*T*) + Γ_*i,T*_ (*s*),

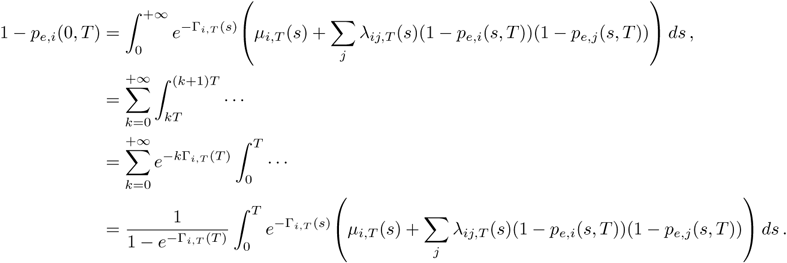

With the change of variables *t* = *s*/*T*, we get

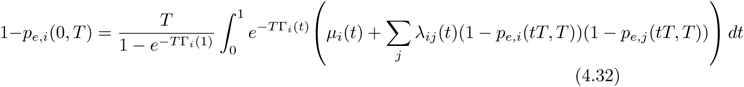

Letting *T* = *T*_*n*_, we can take limits, and we recognise a Laplace integral

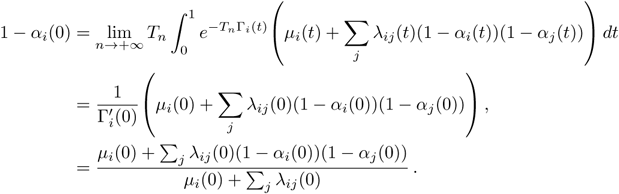

These are exactly the equations satisfied by π_*e,i*_(0). □

### 5 A vector borne disease: Zika Virus

We want to determine the probability of emergence of Zika Virus, a newly emerging vector borne disease of humans. Our starting point is the epidemioloical model of Zika used in previous studies Lourenço et al. [2017], Suparit et al. [2018], Zhang et al. [2017]. The epidemiological dynamics in the human population is described by an SEIR model: susceptible individuals *S*^*H*^, exposed indivuals *E*^*H*^, infected individuals *I*^*H*^ and recovered/removed individuals *R*^*H*^. The epidemiological dynamics in the vector population is described by and SEI model with compartments *S*^*V*^, *E*^*V*^, *I*^*V*^. This yields the following deterministic dynamics:

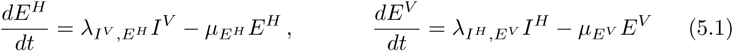

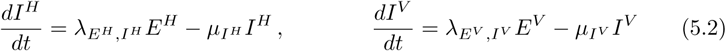

The probability of a major outbreak is the probability of emergence in a birth death process with 4 types. By labelling the states *E*^*H*^, *I*^*H*^, *E*^*V*^, *I*^*V*^ as 1, 2, 3, 4 we can write this system as

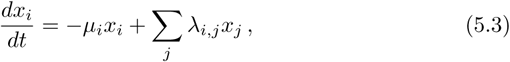

and the generator of the stochastic process is (4.1). Note that we are in a *cyclic* case: *λ*_*i,j*_ = 0 only if *j* = *i* + 1 *mod*(*d*). The death rates *µ*_*i*_ are functions of mortality and recovery rates. For instance, 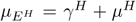 where 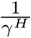 is the expected incubation time of the infection in humans, usually between 5 and 9 days, and 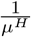 is the expected lifespan of humans, generally taken to be 75 years. The total number of humans *N*^*H*^ = *S*^*H*^ + *E*^*H*^ + *I*^*H*^ + *R*^*H*^ is assumed to be constant over time. Crucially, the seasonality, the periodicity in time, of some parameters comes only through their dependance on temperature. More precisely, a parameter *µ*_*i*_, *λ*_*i,j*_ is periodic iff it is a function of temperature *T*, which is periodic (with period one year). The parameter values infered from observed epidemiological dynamics agree very well among the three studies we used Lourenço et al. [2017], Suparit et al. [2018], Zhang et al. [2017]. All the parameter values used in our models are summed up in Table 1.

**Table 1:** Definition and values of the parameters used in the Zika models. All the rates (*λ, µ* and *b*) are in (*days*)^−1^.

|  | Definition | Model I | Model II |
| --- | --- | --- | --- |
| $\lambda_{E^H, I^H}$ | Rate of transition between $E^H$ and $I^H$ | 0.143 | |
| $\lambda_{I^H, E^V}$ | Rate of transition between $I^H$ and $E^V$ | $bM(T^\circ)$ | $bM(T^\circ)\beta(T^\circ)$ |
| $\lambda_{E^V, I^V}$ | Rate of transition between $E^V$ and $I^V$ | 0.125 | |
| $\lambda_{I^V, E^H}$ | Rate of transition between $I^V$ and $E^H$ | $b$ | $b\beta(T^\circ)$ |
| $b$ | Biting rate of the vector | 0.32 | |
| $\mu_{E^H}$ | Recovery rate + mortality rate of $E^H$ | 0.143 | |
| $\mu_{I^H}$ | Recovery rate + mortality rate of $I^H$ | 0.2 | |
| $\mu_{E^V}$ | Mortality rate of $E^V$ | 0.525 | |
| $\mu_{I^V}$ | Mortality rate of $I^V$ | 0.4 | |
| $T$ | Period of the fluctuation | 365 days | |
| $T_{moy}^\circ$ | Mean temperature | 27° and 29° | 25° |
| $T_{opt}^\circ$ | Optimal temperature for the reproduction of the vector | 32° | |
| $A_{T^\circ}$ | Amplitude of the variation in temperature | 4° | 8° |
| $M_{max}$ | Maximal value of the number of vectors per human $\frac{N^V}{N^H}$ | 10 | |
| $\sigma_{T^\circ}^2$ | Variance for the distribution of the number of vectors per human around the optimal temperature $T_{opt}^\circ$ | 14.05 | |

The seasonality comes through temperature only, which follows a sinusoid:

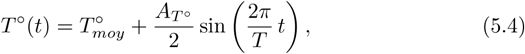

with 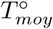 the mean temperature and 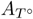 the amplitude of the variation in temperature.

The number of vectors per human 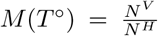 is assumed to reach a maximum *M*_*max*_ at the optimal temperature 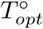. Following Zhang et al. [2017] we assume the number of vectors per human drops away from the optimal temperature:

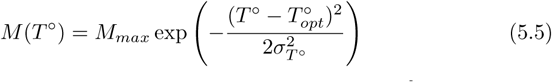

with the optimal temperature for vector reproduction and 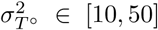 a spread factor (the “variance” for this gaussian shape).

In this model the *winter* (i.e. the low transmission season) is defined to be the set of times *t*_0_ for which the computation of *R*_0_ would yield *R*_0_ *<* 1 if the parameters were to be frozen at their values *λ*_*ij*_(*t*_0_) and *µ*_*i*_(*t*_0_). In other words it is the set of time for which the product of the birth rates is less than the product of the death rates.

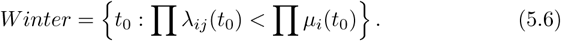

Unfortunately, unlike the model in dimension 1, where it is defined as the complementary of the set *A* of Proposition 2.1, the timing of the *winter is coming* effect cannot be identified easily (see Figure S1). We decided to define the *winter is coming* time zone to be the set of times for which the emergence probability is less than a fixed threshold *δ* = 0.05

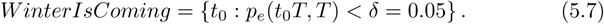

#### Model I

In this simple model, as in Suparit et al. [2018], there is a single rate that varies with time. We assume that seasonality affects only the population size of vectors and, consequently, the only rate that varies with time is 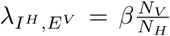. In the figure 5 of the main text we show the influence of a small shift in the mean temperature on the probability of pathogen emergence after the introduction of a single infected human in the *exposed* state. This figure illustrates that we recover the winter is coming effect when the amplitude of the seasonal variation are large enough.

In the figure S5 we illustrate the influence of the state of the introduced pathogen: *E*^*H*^, *I*^*H*^, *E*^*V*^ or *I*^*V*^. This figure shows that the probability of emergence is much higher when the pathogen is introduced in a human host (black lines) rather than in a vector (red lines) because the duration of infection is higher in humans. Besides, the probability of emergence is higher if the introduced host is already infectious (dashed line) rather than exposed (full line) because the pathogen can readily be transmitted from an infectious host and does not need to wait the end of the incubation time.

**Figure S5:**
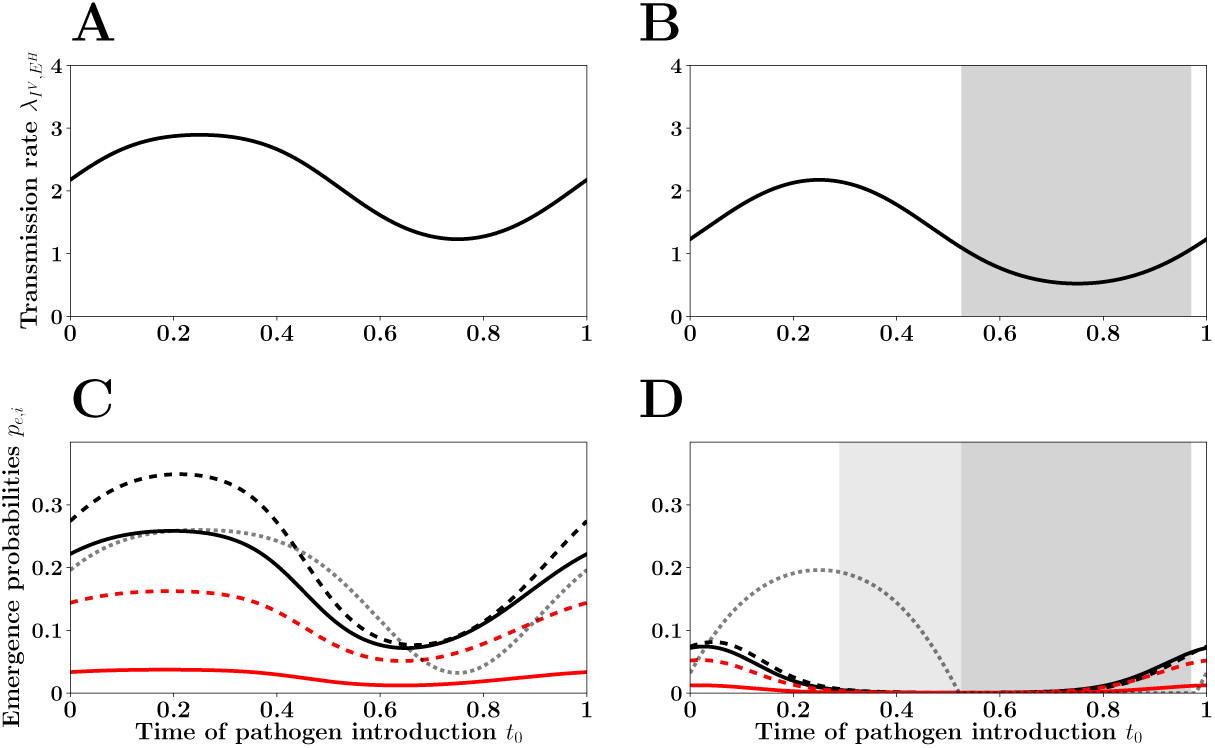
Effect of the state of the introduced pathogen on the probability of Zika emergence (model I). Same as figure 5 in the main text but here we illustrate the effect of the state in which the pathogen is introduced (in figures C and D): in solid black an exposed human (*E*^*H*^), in dashed black an infected human (*I*^*H*^), in solid red an exposed mosquito (*E*^*V*^), in dashed red an infected mosquito (*I*^*V*^).

#### Model II

In this second model we allow more transition rates to depend on the temperature as in Zhang et al. [2017] and Lourenço et al. [2017]. We chose to have two transmission parameters depending on the temperature 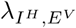 and 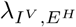. They both have a common factor, the *biting rate*, modeled after Lambrechts et al. [2011] as:

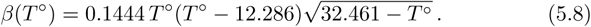

This yields:

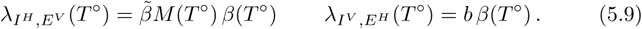

Figure S6 shows the fluctuations in both 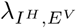 and 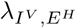 throughout the season. The function *β*(*T°*) induces major drops in transmission when the temperature is too low but also when it is too high relative to 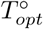. These fluctuations induce two peaks in the probability of pathogene emergence. The optimal timing of the control can have a massive impact on the risk of pathogen emergence during the period of control but also before. This confirms the relevance of the *winter is coming* effect in a realistic model of Zika emergence.

**Figure S6:**
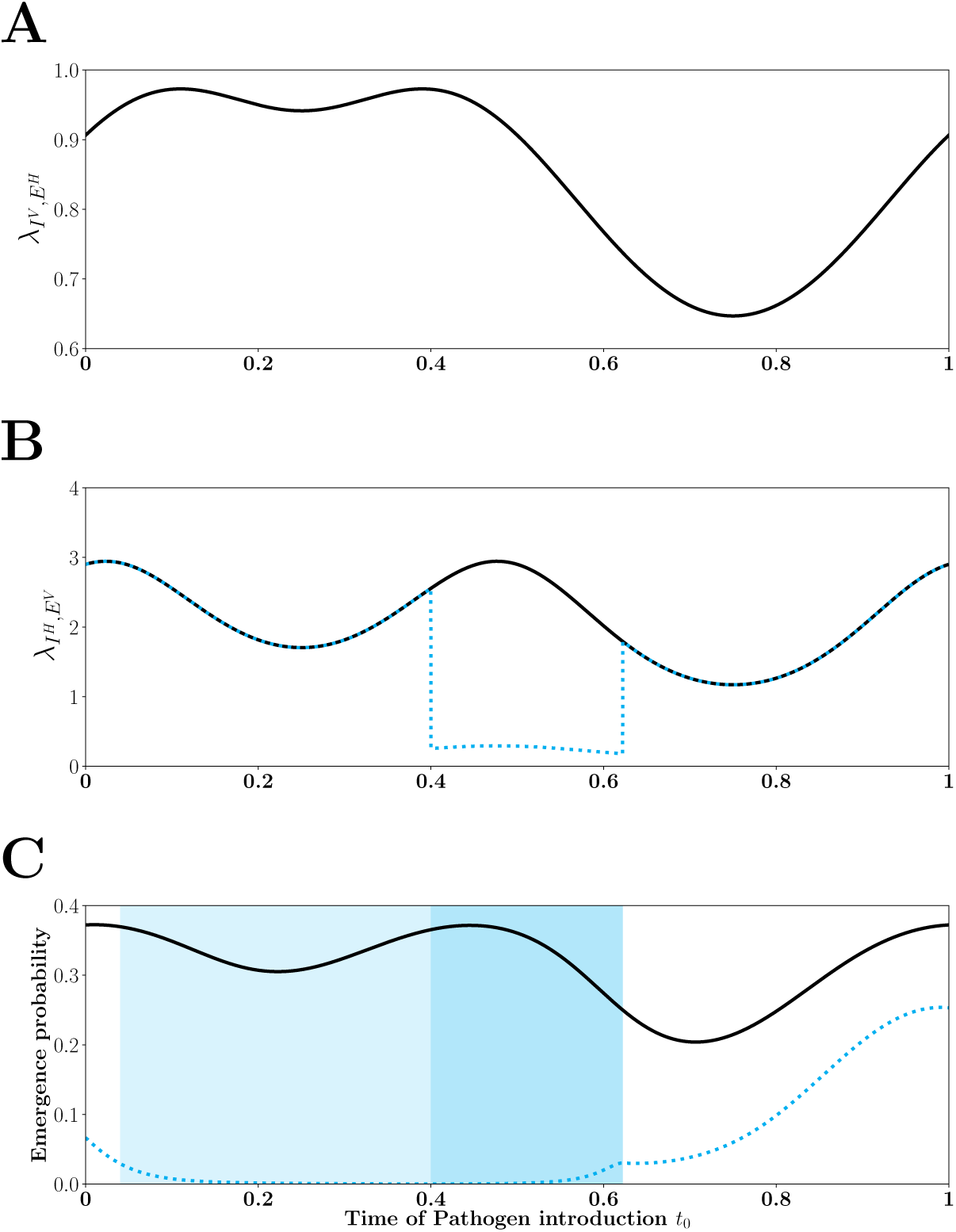
Probability of Zika emergence (model II). In model II both the density of mosquito vectors (equation 5.5)and the biting rate (equations 5.8 and 5.9) fluctuate throughout the season. This yields a fluctuation in 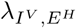 (A) and 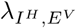 (B). Without control the probability of Zika emergence (in solid black) is always above 0.2 because transmission remains relatively high in the low-transmission season (see also figure 5A and 5C). The optimal control operates only on the density of mosquitoes, the blue dotted curve in B, during the time interval [*t*_1_, *t*_2_] = [0.4, 0.62] (blue shading in C) and results in an important *winter is coming* effect (light blue shading in C).

### 6 Pulse interventions

#### 6.1 A pulse vaccination model

We want to determine the probability of pathogen emergence in a seasonal environment when vaccination is applied periodically (every year) in a pulse occuring at time *δ* during the year. The question is: *When is the best time δ to vaccinate?* Following Grassly and Fraser [2006] we use a one dimensional linear birth and death process with a constant death rate *µ*(*t*) = 1 and a time varying birth rate:

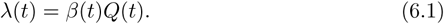

The function *β*(*t*) refers to the seasonal transmission rate:

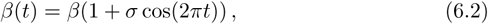

The function *Q*(*t*) is the proportion of unvaccinated in a deterministic model of population with death rate and birth rate equal to *µ*, with *Q*(*t*) + *V* (*t*) = 1 and thus for *t* ∈ [0, 1):

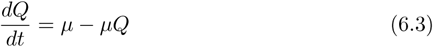

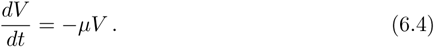

The solution of this ODE is:

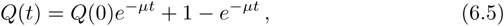

Let us assume that a pulse vaccination is applied at the end of the season (at time 1). The number of unvaccinated is instantly decreased by a factor *q* ∈ (0, 1):

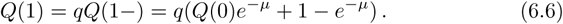

Then we make the same operation on intervals [1, 2[, [2, 3[, *…*. It is easy to see that the function constructed this way is asymptotically close to the 1-periodic function:

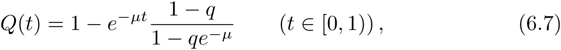

which is the function (4.4) of Grassly and Fraser [2006].

Next we assume that the pulse vaccination may be applied at another point in time during the year. The parameter *δ* ∈ (0, 1) refers to the shift between the seasonal fluctuation and the vaccination pulse. We consider *Q*_*δ*_(*t*) = *Q*(*t* − *δ*) and *λ*_*δ*_(*t*) = *Q*_*δ*_(*t*)*β*(*t*). We obtain

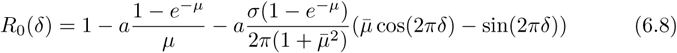

with 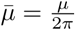 and a 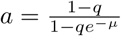.

**Figure S7:**
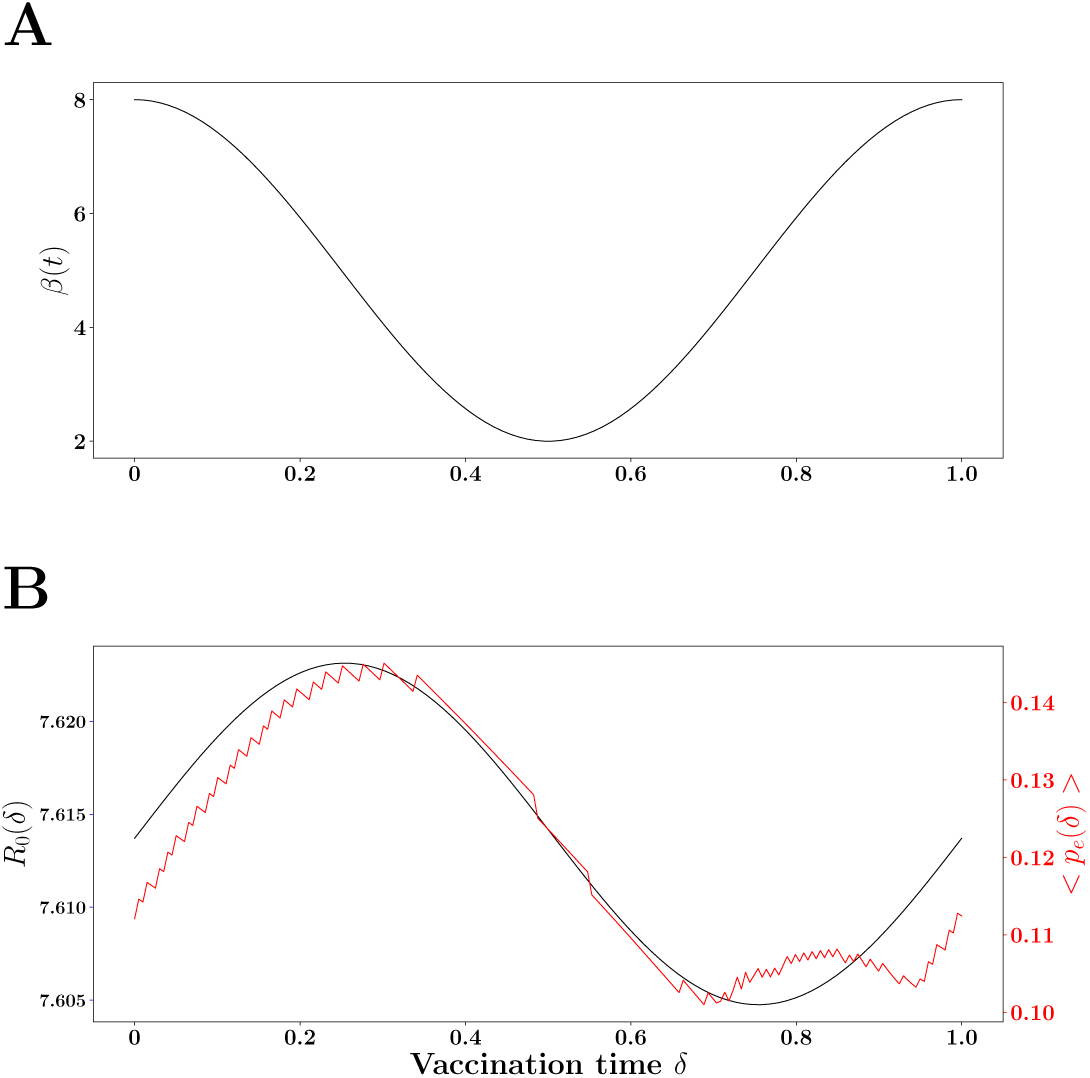
Effect of the timing of the pulse vaccination on *R*_0_ and the mean probability of pathogen emergence. The upper curve is *β*(*t*), pathogen transmission rate before vaccination (figure A): it enables to identify the high and low transmission seasons. The mean emergence probability without pulse vaccination is *⟨p*_*e*_*⟩* = 0.75. In figure B we plot *R*_0_(*δ*), the basic reproduction number (black line), and *< p*_*e*_(*δ*) *>*, the mean extinction probability (red line) against *δ*, the timing of the pulse vaccination (i.e. the amount of time after the start of the high transmission period). Parameters: *σ* = 0.6, *β* = 5.0, *q* = 0.6, *µ* = 0.16.

The optimal shift,that minimizes *R*_0_(*δ*), has been computed by Grassly and Fraser [2006] and is 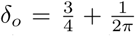 arctan 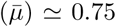 for small *µ* (say *µ* = 0.16). This result is in accordance with Grassly and Fraser [2006]. They found by doing some numerical stochastic simulations that *δ* = 0.75 was a better choice than *δ* = 0.5 to decrease the mean emergence probability *< p*_*e*_ *>*. Our calculation, however, allows to compute precisely the optimal *δ* using the original formula of Kendall [1948] adapted to seasonal environment, equation (1.3). We find that the optimal delta *δ*_*\**_, that minimizes the mean emergence proability *⟨p*_*e*_*⟩*, for the set of parameters given by Grassly and Fraser [2006], *δ*_*\**_ = 0.69, see figure S7. Again, we see that the control strategy that minimizes *⟨p*_*e*_*⟩* is not the same as the strategy that minimizes *R*_0_.

#### 6.2 A pulse treatment model

We want to determine the probability of pathogen emergence in a seasonal environment when an antibiotic treatment is applied periodically (every year) in a pulse occuring at time *δ* during the year. The question is: *When is the best time δ to treat?*

In the absence of treatment we use the same one dimensional model where the death rate *µ*(*t*) is constant and the birth rate is:

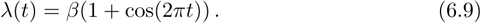

The treatment pulse occurs at time *δ* ∈ (0, 1): there is an increase of *µ*(*t*) from 1 to 1 + Δ*µ* during the time interval (*δ, δ* + Δ*t*). If Δ*t* is small enough, this is close to a dirac pulse, and the effect is to reduce quasi instantly the number of infected by the factor *c* = *e*^−Δ*t*Δ*µ*^. Since *µ*_*δ*_(*t*) = 1 + Δ*µ* **1**_(*δ<t<t*+Δ*t*)_, the basic reproduction number is given by:

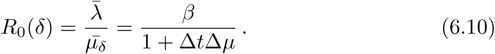

Observe that *R*_0_(*δ*) *does depend* on the pulse intensity Δ*t*Δ*µ*, but *does not depend* on the timing *δ* of the pulse.

Nevertheless, the emergence probability *does depend* on *δ*, and we can determine the optimal *δ*_*\**_ that minimizes the mean emergence probability *< p*_*e*_(*δ*) *>* see Figure S8. This scenario illustrates again that minimizing *R*_0_(*δ*) or minimizing *< p*_*e*_(*δ*) *>* does not yield the same optimal control strategy. We also recover that the optimal timing of the control is just before the high transmission season (compare S7 and S8).

**Figure S8:**
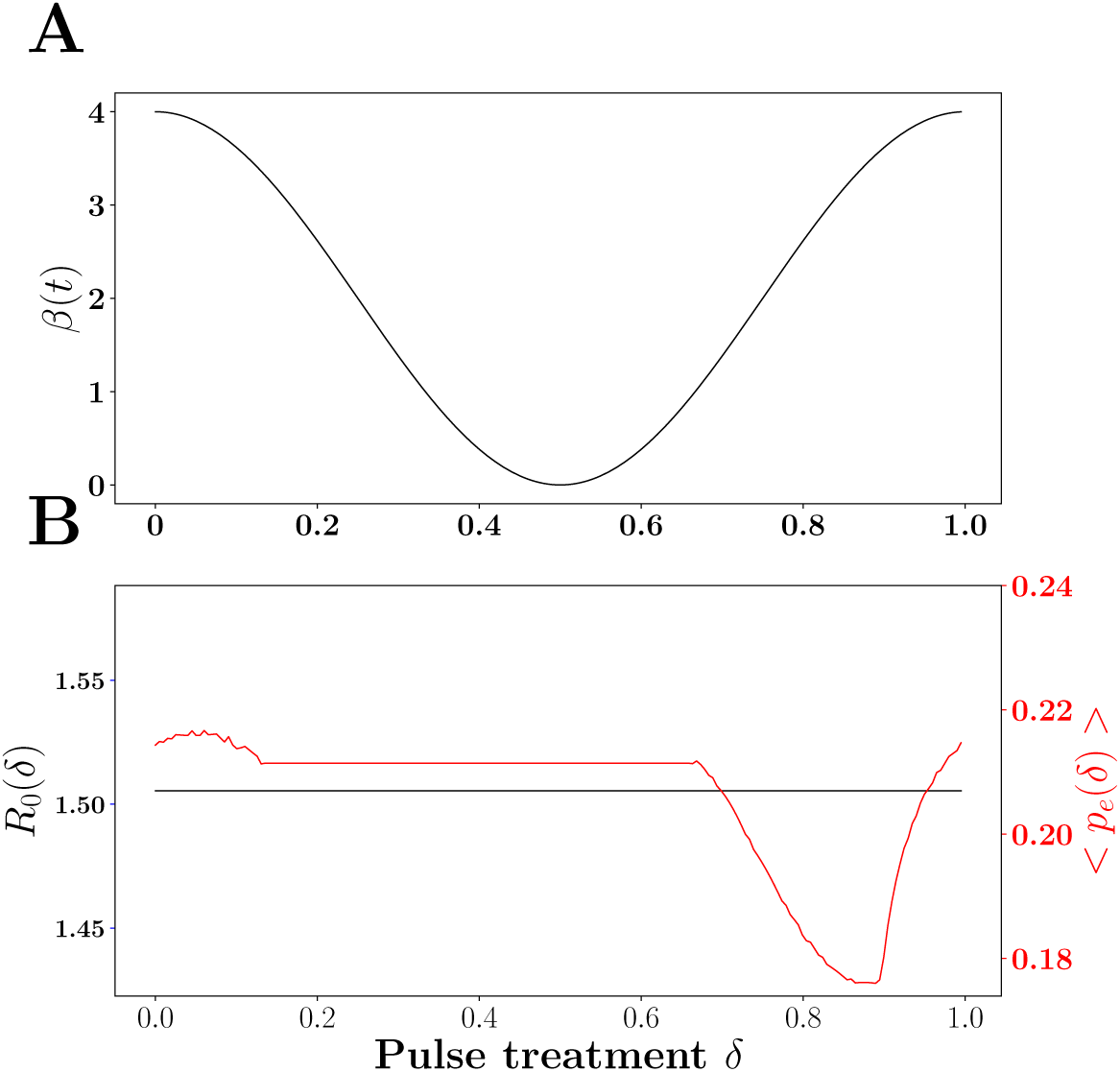
Effect of the timing of the pulse treatment on *R*_0_ and the mean probability of pathogen emergence. The upper curve is *β*(*t*), pathogen trasnsmission rate before control (figure A): it enables to identify the high and low transmission seasons. When no treatment is used the basic reproduction ratio is *R*_0_ = 1.505 and the mean emergence probability is *⟨p*_*e*_*⟩* = 0.302. In figure B we plot *R*_0_(*δ*), the basic reproduction number (black line), and *< p*_*e*_(*δ*) *>*, the mean extinction probability (red line) against *δ*, the timing of the pulse treatment (i.e. the amount of time after the start of the high transmission period). Parameters: *β* = 2.0, Δ*t* = 0.01, Δ*µ* = 32.

### 7 A density dependent example

The above analysis relies on the branching process assumption where pathogen growth is not affected by density dependent effects. Yet, after some time, the spread of the pathogen is likely to reduce the density of susceptible hosts and this will feed back on the malthusian growth rate of the pathogen population. For instance, let us consider the following epidemiological model:

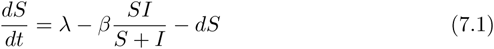

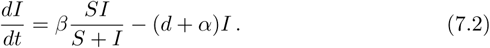

Where *λ* is the rate at which new susceptible hosts enter the population per unit area, *d* is natural host death rate, *α* is the additional mortality induced by virulence and *β* is pathogen transmission rate. The above equations can be viewed as the deterministic limit of a continuous time Markov chain model that tracks the dynamics of *S*^(*n*)^, the finite population size of susceptible hosts, and *I*^(*n*)^, the finite population size of infected hosts. The deterministic model is the limit for *n*→ ∞ of the stochastic model for *S*^(*n*)^*/n*, the density of the population of susceptible hosts, and *I*^(*n*)^*/n*, the density of the population of infected hosts (see for instance Parsons et al (2018) for a derivation). When all the rates are assumed to be constant the equilibrium finite size of the population of infected hosts is equal to:

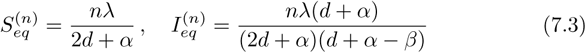

In the following, however, all the rates are assumed constant, except *β* which is *T* -periodic:

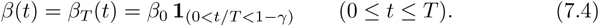

The associated branching process has periodic birth rate *λ*(*t*) = *β*_*T*_ (*t*) and constant death rate *µ* = *d* + *α*. To study the effect of density dependence we performed individual based simulations to obtain the probability of emergence (the ability of the pathogen population to survive after 10 periods) after the introduction of a single infected host at time *t*_0_. Figure S9 illustrates the good match between the stochastic simulations and the computation of *p*_*e*_(*t*_0_*T, T*) under the assumption that transmission follows a square wave.

**Figure S9:**
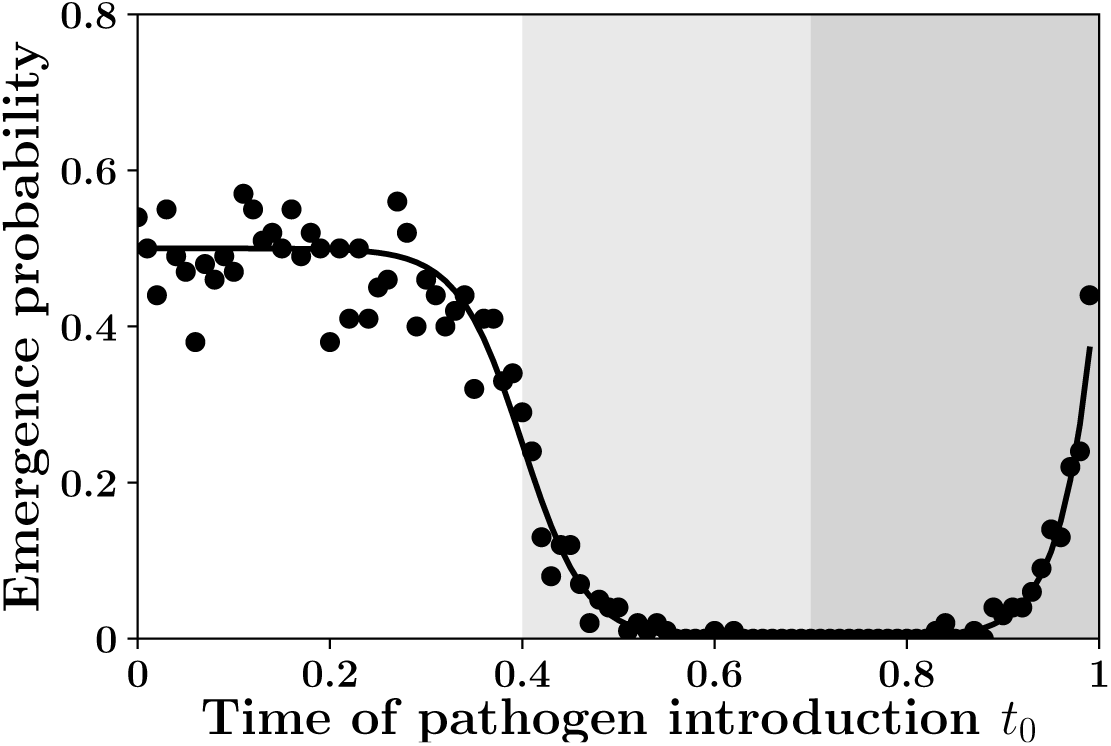
Individual based simulations and numerical derivations of the probability of pathogen emergence. We found a good match between stochastic simulations (each black dot is obtained from 1000 replicates) and numerical derivations of the probability *p*_*e*_(*t*_0_*T, T*) of pathogen emergence (black line). As in figures 2 and S1 the low transmission period is indicated by the gray shading and the *winter is coming* effect by the light gray shading. Parameters: *β*_0_ = 0.2, *d* = 0.09, *α* = 0.01, *γ* = 0.3, *λ* = 1000, *n* = 10, *T* = 300.

The absence of effect of density dependence in figure S9 is due to the large size of the population of infected hosts. Indeed, for the parameter values used in figure S9 the size of the population of infected hosts is expected to be around 52630 in the high transmission season (see equation 7.3). However, when the size of the population of infected hosts drops, the risk of pathogen extinction during the low transmission period increases. Consequently, the *winter is coming* effect is increased by density dependence. This effect can be very high and can lead the probability of pathogen emergence to zero on the whole period (see figure S10).

In the following we use an heuristic argument to explain the effect of finite population size on pathogen emergence in seasonal environments. Let us focus on the the probability of emergence *p*_*e*_(0, *T*) for a pathogen introduced at time 0. If it escapes early extinction the pathogen population will grow at a rate *r*(*t*) *>* 0 in the high transmission season. If the system size is sufficiently large, by the law of large numbers, the stochastic dynamics of *I*^(*n*)^(*t*)*/n* is very close to the deterministic solution of 7.1. So let us assume, for our heuristic, that if the infected pathogen introced at time 0 survives, at time *T* (1 − *γ*) there are exactly 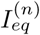 infected hosts (see equation 7.3).

During the low transmission period [*T* (1 − *γ*), *T*] the number of infected hosts is a death process: infected hosts die with constant rate *µ* = *d* + *α*. Therefore, the probability that an infected survives the winter, of length *γT* is given by 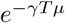 and the probability that the 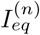 infected hosts survive the winter is:

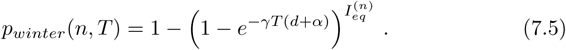

Hence, our heuristic gives the upper bound, *P*_*e*_(0, *T*) *≤ p*_*e*_(0, *T*)*p*_*winter*_(*n, T*)^*W*^ where *W* is the number of winters we consider, and *p*_*e*_(0, *T*) is the probability of emergence starting from zero for the associated branching process.

The curve *n* → *p*_*winter*_(*n, T*)^*W*^ is close to a step function: close to 0 before *n*_*c*_, and close to 1 after *n*_*c*_, where *n*_*c*_ is the critical area size of the system defined by the equation: *p*_*winter*_(*n*_*C*_, *T*) = 1*/*2 and equals:

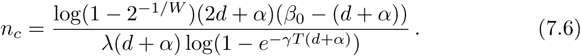

Figure S10 shows results from individual based simulations (black dots) that confirm the validity of our heuristics: when *n < n*_*c*_ the probability of pathogen emergence is droping to very low levels, and when *n > n*_*c*_ the probability of pathogen emergence is close to *p*_*e*_(0, *T*) computed for the branching process (i.e. without density dependence).

**Figure S10:**
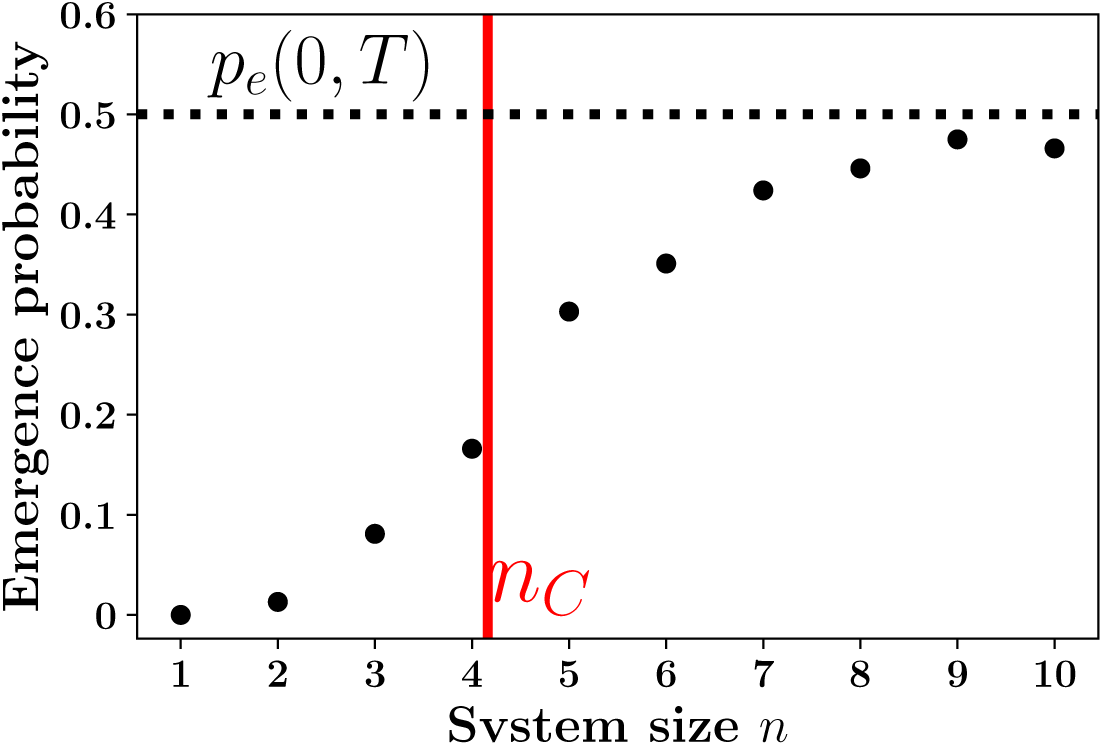
Individual based simulations and numerical derivations of the probability of pathogen emergence *p*_*e*_(0, *T*) against the system size. *n*. We used same parameter values as in figure S9 but we varied the size of the system size between *n* = 1 and 10. Our approximation for the critical system size *n*_*c*_ = 4.16 matches the point at which the probability of pathogen emergence drops. Note that our branching process approximation of the probability of pathogen emergence (i.e. *p*_*e*_ = 0.5) matches results of individual simulation when *n ≫ n*_*c*_. Parameters: *β*_0_ = 0.2, *d* = 0.09, *α* = 0.01, *γ* = 0.3, *λ* = 1000, *n* = 10, *T* = 300.

### 8 Additional computations and proofs

#### Proof of Proposition 3.1

Remember that we assumed that 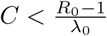, that is *φ*_*ρ*_(1) *>* 0. Forgetting about the limiting case, which we obtain by having equalities instead of inequalities, we have six cases to consider see figure S11.

**Figure S11:**
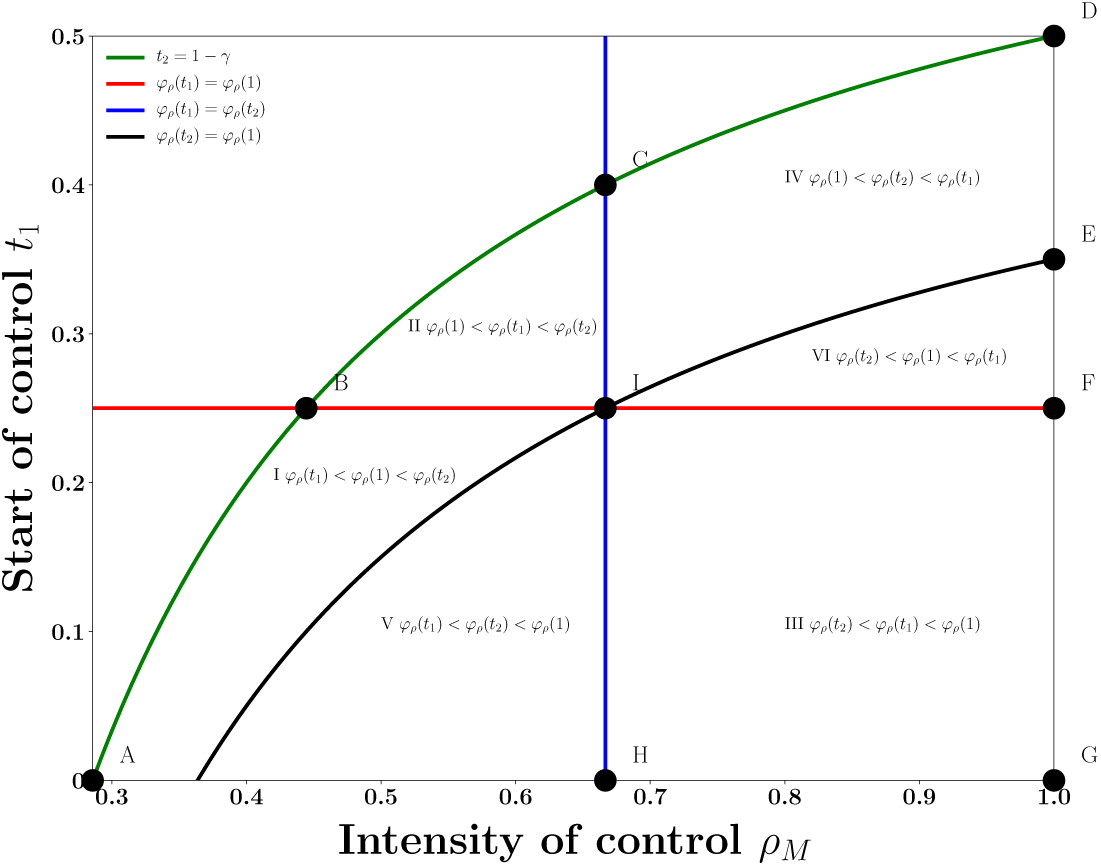
Analysis of figure 4. We plot the six different cases (see proof of Proposition 3.1) corresponding to the respective positions of the three numbers *φ*_*ρ*_(*t*_1_), *φ*_*ρ*_(*t*_2_), *φ*_*ρ*_(1). Zones II,III,IV and VI correspond to an optimal control. The dashed red contour in figure 4 corresponds exactly to the area delineated by points B,C,D,E,F,G,H,I and B.)

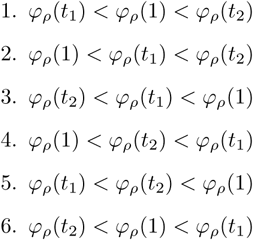

The different shapes of the integrated rate function are depicted in figure S12. This figure is a companion to the proof below, since one can use this figure to spot the local minima and the corresponding *t*^*\**^ times using the geometric construction presented in figure S1.

Finally, the actual controlled rates corresponding to the particular points of figure S11 are given in figure S13.

**Figure S12:**
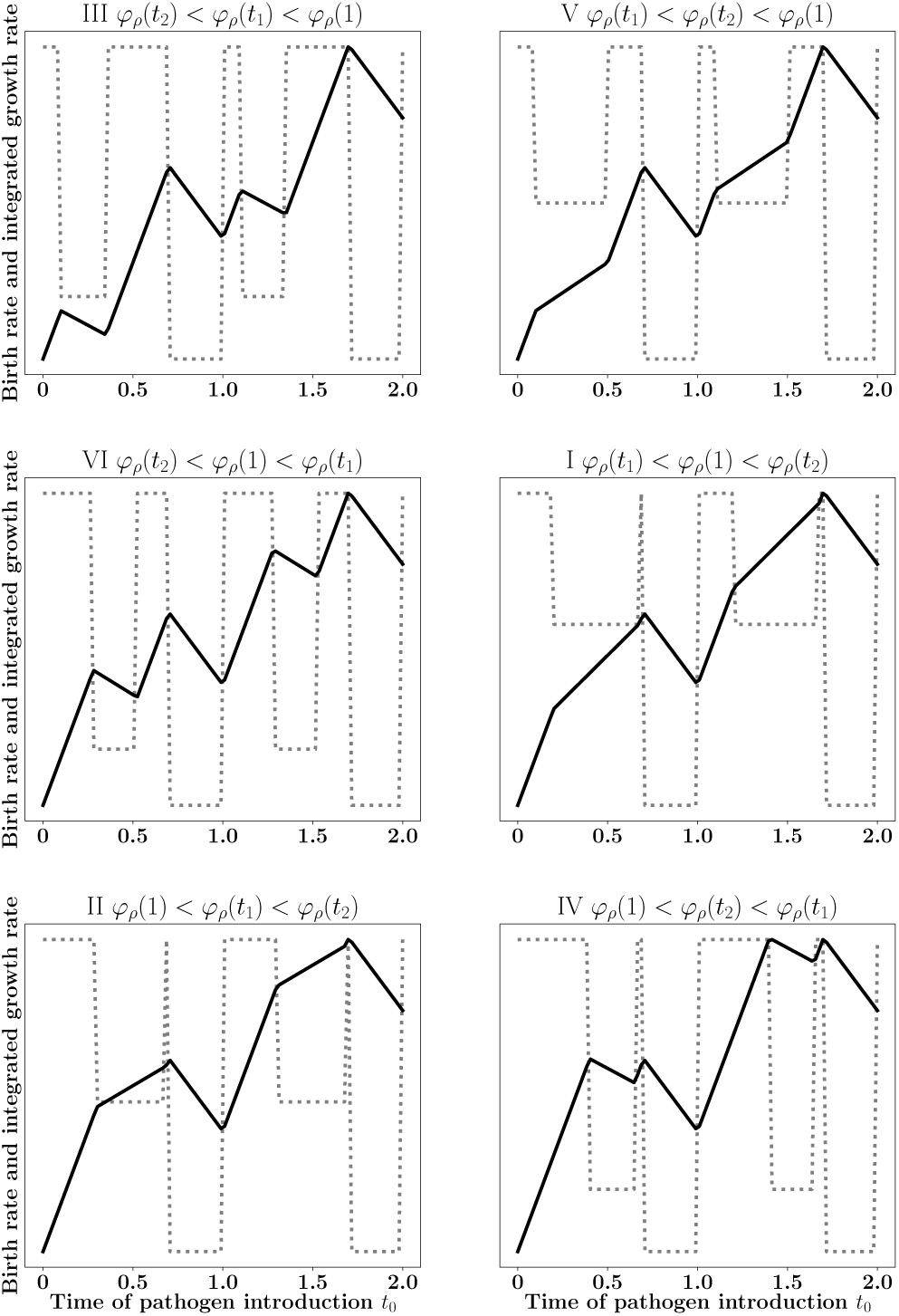
Understanding the different cases in figure S11. The dotted black line is the controlled birth rate *λ*_*ρ*_(*t*), the solid black line is the controlled integrated growth rate *φ*_*ρ*_(*t*).

**Figure S13:**
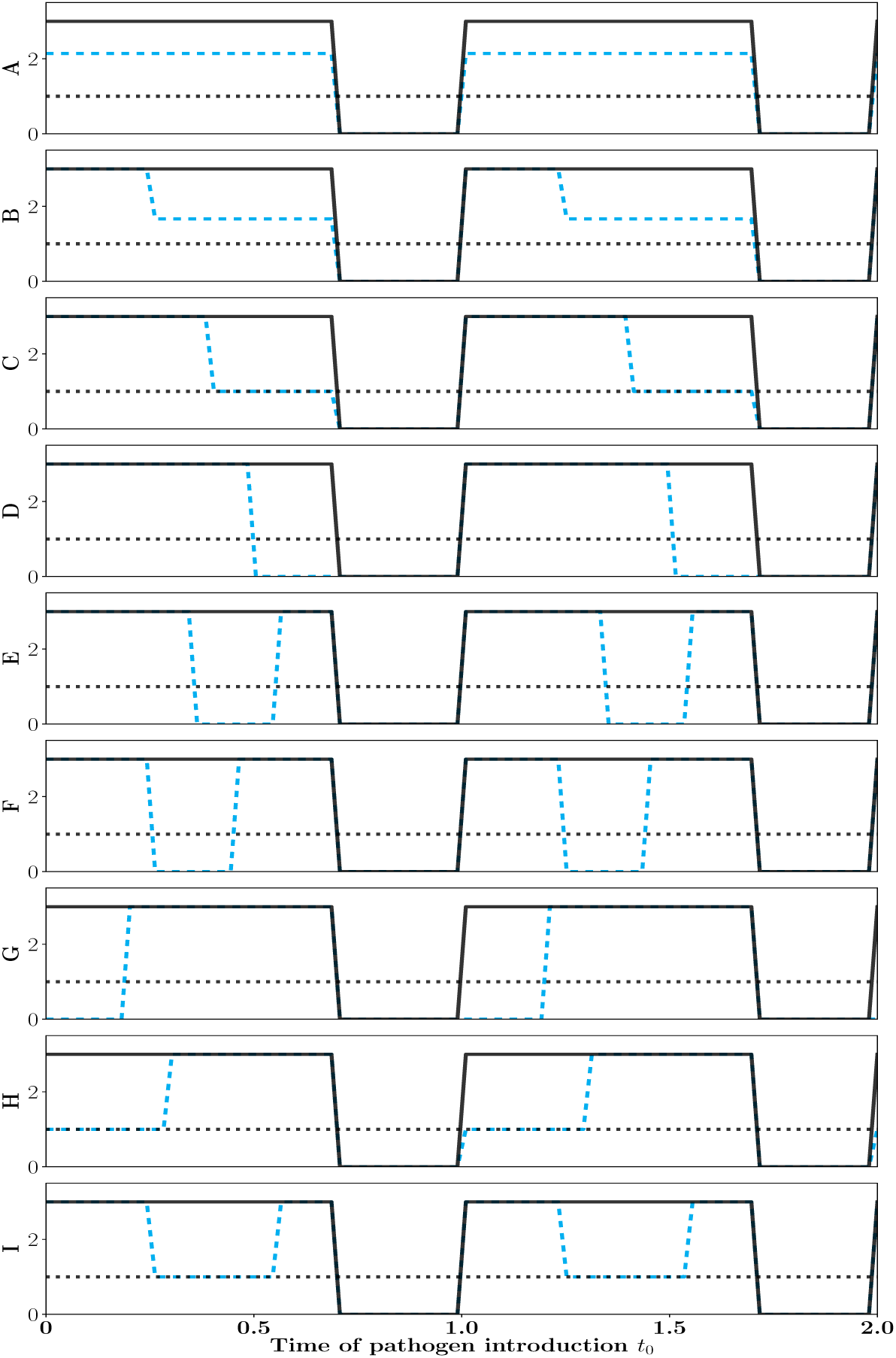
Understanding the 9 different points in figure S11. The solid black line is the uncontrolled birth rate *λ*(*t*) = *λ*_0_ **1**_(0*<t<*1−*γ*)_, the dashed blue line is the controlled birth rate *λ*_*ρ*_(*t*), the dotted black line is the death rate *µ* = 1. Parameters are *λ*_0_ = 3.0, *γ* = 0.3, *C* = 0.2.

**Case I**: *φ*_*ρ*_(*t*_1_) *< φ*_*ρ*_(1) *< φ*_*ρ*_(*t*_2_). Then we have 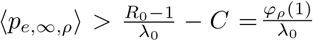. the strategy is not optimal.

Indeed let 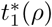 be the unique *t* ∈ (*t*_1_, *t*_2_) such that 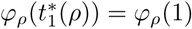. Then

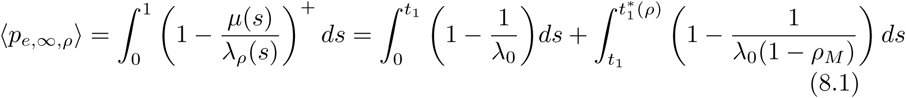

Since we have

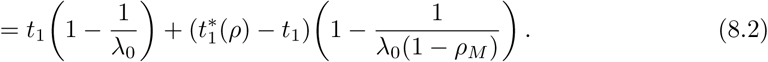

we get that

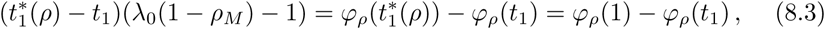

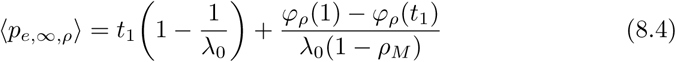

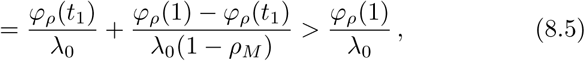

since *φ*_*ρ*_(1) *> φ*_*ρ*_(*t*_1_).

**Case II**: *φ*_*ρ*_(1) *< φ*_*ρ*_(*t*_1_) *< φ*_*ρ*_(*t*_2_). Then we have 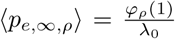: the strategy is optimal.

Indeed let 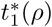 be the unique *t* ∈ (0, *t*_1_) such that 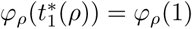. Then

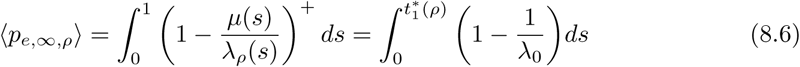

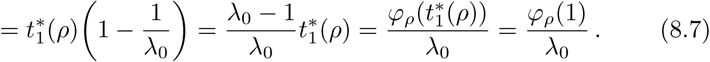

**Case III**: *φ*_*ρ*_(*t*_2_) *< φ*_*ρ*_(*t*_1_) *< φ*_*ρ*_(1). This is also an optimal case: 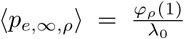. We let 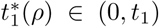 such that 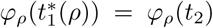 and 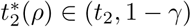 such that 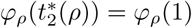. Then

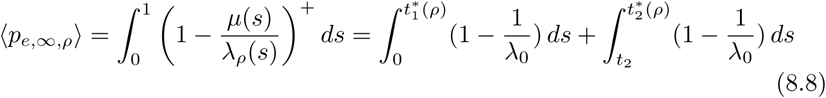

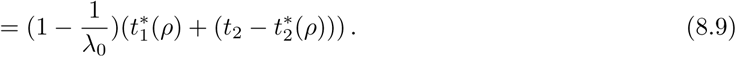

Since we have

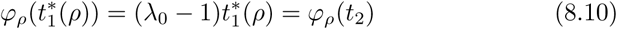

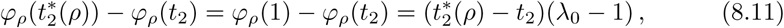

we get that

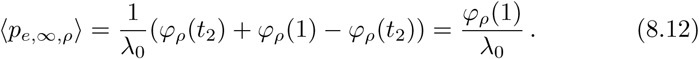

**Case IV:** *φ*_*ρ*_(1) *< φ*_*ρ*_(*t*_2_) *< φ*_*ρ*_(*t*_1_). This is also an optimal case: 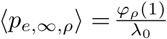. We let 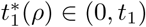 such that 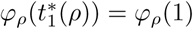 and we get

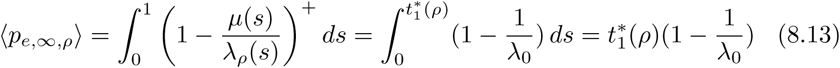

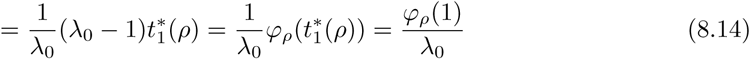

**Case V**:*φ*_*ρ*_(*t*_1_) *< φ*_*ρ*_(*t*_2_) *< φ*_*ρ*_(1). The strategy is not optimal. Let 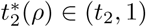 such that 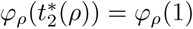. Then

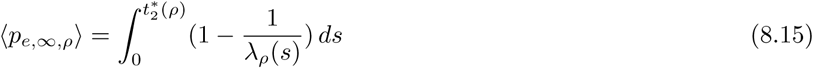

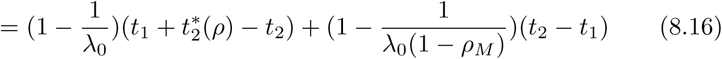

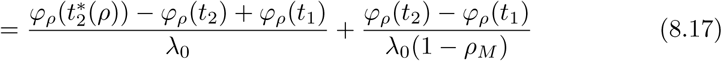

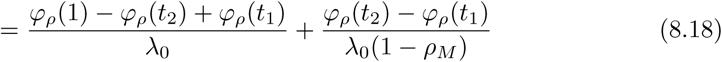

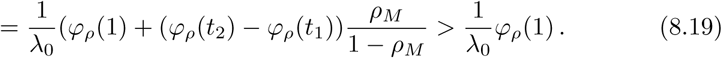

**Case VI:** *φ*_*ρ*_(*t*_2_) *< φ*_*ρ*_(1) *< φ*_*ρ*_(*t*_1_). This is an optimal strategy. Let 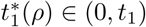 such that 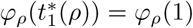.Then

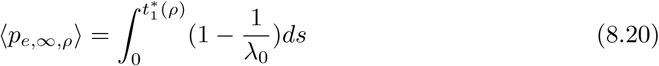

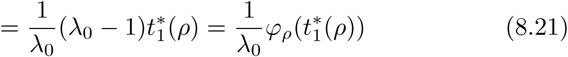

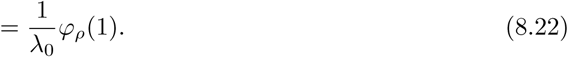

